# Aging hastens locomotor decline in PINK1 knockout rats in association with decreased nigral, but not striatal, dopamine and tyrosine hydroxylase expression

**DOI:** 10.1101/2024.02.01.578317

**Authors:** Isabel Soto, Robert McManus, Walter Navarrete-Barahona, Ella A. Kasanga, Kirby Doshier, Vicki A. Nejtek, Michael F. Salvatore

**Affiliations:** Department of Pharmacology & Neuroscience, University of North Texas Health Science Center, Fort Worth, TX 76107

**Author notes:** **Corresponding author:** Michael F. Salvatore, PhD. Department of Pharmacology & Neuroscience University of North Texas Health Science Center 3500 Camp Bowie Blvd Fort Worth, Texas 76107.

**Keywords:** Substantia nigra, Parkinson’s disease, PINK1, mitochondria, animal models, aging

## Abstract

Parkinson’s disease (PD) rodent models provide insight into the relationship between nigrostriatal dopamine (DA) signaling and locomotor function. Although toxin-based rat models produce frank nigrostriatal neuron loss and eventual motor decline characteristic of PD, the rapid nature of neuronal loss may not adequately translate premotor traits, such as cognitive decline. Unfortunately, rodent genetic PD models, like the Pink1 knockout (KO) rat, often fail to replicate the differential severity of striatal DA and tyrosine hydroxylase (TH) loss, and a bradykinetic phenotype, reminiscent of human PD. To elucidate this inconsistency, we evaluated aging as a progression factor in the timing of motor and non-motor cognitive impairments. Male PINK1 KO and age-matched wild type (WT) rats were evaluated in a longitudinal study from 3 to 16 months old in one cohort, and in a cross-sectional study of young adult (6-7 months) and aged (18-19 months) in another cohort. Young adult PINK1 KO rats exhibited hyperkinetic behavior associated with elevated DA and TH in the substantia nigra (SN), which decreased therein, but not striatum, in the aged KO rats. Additionally, norepinephrine levels decreased in aged KO rats in the prefrontal cortex (PFC), paired with a higher DA content in young and aged KO. Although a younger age of onset characterizes familial forms of PD, our results underscore the critical need to consider age-related factors. Moreover, the results indicate that compensatory mechanisms may exist to preserve locomotor function, evidenced by increased DA in the SN early in the lifespan, in response to deficient PINK1 function, which declines with aging and the onset of motor impairment.

## Introduction

Parkinson’s disease (PD) is a multifactorial neurodegenerative disease with aging as the leading risk factor (Gelders et. al, 2018; Hall 2018 et al.; Hindle et al., 2010; Collier et al., 2011; 2017). Nearly one million people in the USA are currently diagnosed with PD, and this number is rapidly rising alongside the aging population (de Lau et al., 2016; Marras et al., 2018). Characterized by a loss of dopaminergic (DA) neurons from the substantia nigra (SN) and abnormal alpha-synuclein aggregation, PD manifests with symptoms such as bradykinesia, muscle rigidity, tremors, and gait impairments (Stefanis et al., 2012; Stoker et al., 2018). Additional non-motor symptoms (NMS), such as early decline in executive functioning, stems from impaired norepinephrine (NE) and DA signaling in the prefrontal cortex (PFC) that can be observed in up to one-third of patients (Aarsland et al., 2017; Braak et al., 2003; Postuma et al., 2019, Nejtek et al., 2021, Durcan et al., 2019; Dirnberger et al., 2013; Salvatore et al., 2021). Notably, the motor symptoms of PD arise years into the disease course, and therefore patients are often pathologically well advanced prior to seeking medical attention, which hinders the ability to assess the earliest neurobiological processes of the disease (Berg et al., 2021; Postuma et al., 2019; Kordower et al., 2013; Hauser et al., 2018).

Rodent PD models may provide mechanistic insights into the stages of neuropathology that align with behavioral traits. However, while aging plays a crucial role in PD development, its neurobiological underpinnings are often overlooked and inadequately translated from animal models (Zeiss et al., 2017; Barker and Björklund, 2020). Moreover, whereas toxin models like 6-hydroxydopamine (6-OHDA) can induce nigrostriatal neuron loss of similar magnitude to that in human PD (Kasanga et al., 2023), they have been continuously criticized for the rapid nature of nigrostriatal neuron loss-presupposing (without evidence) that the rate of progressive loss occurring for years prior to motor symptom onset in human PD is not emulated (Bezard et al., 2013; Dauer et al., 2003). Unfortunately, efforts to mimic the human condition in rodent models with genetic or synuclein-based manipulation have yet to yield a reliable and consistent behavioral phenotype (Polinski, 2021) in concert with the hallmark changes of progressive loss of nigrostriatal neuron terminals and, to a lesser degree, cell body loss consistently seen in human PD (Kordower et al., 2013; Heng et al., 2023; Sun et al., 2013; Zhang et al., 2022).

PTEN-Induced Kinase 1 (PINK1) is a protein involved in mitophagy. Its function is compromised in one of the familial forms of PD, leading to early-onset PD with a median age of onset of 30-39 years old (Khan et al., 2002; Valente et al., 2004; Hatano et al., 2004; Ibanez et al., 2006; Ricciardi et al., 2014; Borsche et al., 2020). Translating the phenotype of this mutation into in an animal model has been problematic. The PINK1 knockout (KO) mouse does not exhibit an early onset parkinsonian phenotype, as hoped in developing this model, or loss of nigrostriatal neurons or dopamine (DA) (Kitada et al., 2007; 2009). Instead, the PINK1 KO rat appears to emulate a slower onset and progression of motor impairment (Dave et al., 2014; Grant et al., 2015; Ferris et al., 2018; Grigoruţă et al., 2020). These less-than-ideal characteristics have led to considerable inconsistency in the timing and severity of the onset of a parkinsonian motor phenotype, beginning as early as 4 months months old (Dave et al., 2014), at 8 months old (de Hass et al.,2019; Grant et al., 2015), or not at all unless an environmental stressor is present (Grigoruta et al., 2020). Notably, as PD also impairs cognitive function, some studies with this model have identified memory and learning impairments in the Pink1 KO rat (Pinizzotto et al., 2022; Maynard et al., 2020; Cai et al., 2019; Hoffmeister et al., 2022; Soto et al., 2024).

The age of onset for humans with a PINK1 mutation is highly variable, ranging from 11 to 67 years of age, due to varying point mutations to large deletions (Bentivoglio et al., 2001; Kasten et al., 2018). Likewise, environmental factors and lifestyle may contribute to heterogeneity in the biological response to a PINK1 mutation (Cazeneuve et al., 2009; Taghavi et al., 2017; Steele et al., 2015), as suggested in the KO rat (Grigoruţă et al., 2020). However, only one study has addressed whether aging may be a relevant factor in the inconsistent timing of motor phenotype onset against impaired DA signaling (Zhi et al., 2019). Thus far, only moderate loss, if any, of nigrostriatal neurons, DA, or tyrosine hydroxylase (TH) has ever been reported in this model (Dave et al., 2014; Villeneuve et al., 2016), and any DA loss is nowhere near the severity seen in human PD at diagnosis (Villeneuve et al., 2017; Dave et al, 2014; Creed et al., 2019, 2020; Zhi et al., 2019; Salvatore et al., 2022). Moreover, striatal DA or TH loss has not correlated with motor impairments (Dave et al., 2014; de Haas et al., 2019; Marquis et al., 2020). As such, we postulated that the impact of Pink1 mutation on locomotor function and DA signaling may be more comparable to human PD with advancing age.

To address if aging affects the nigrostriatal changes in TH and DA regulation in line with motor decline, we conducted a longitudinal study comparing male PINK1 KO and WT rats from 3 to 16 months old, and in another cohort, a cross-sectional study with both young adult (6-7 months) and aged (18-19 months) KO and WT rats. We evaluated both motor and cognitive functioning, and tissue levels of DA, NE, and TH protein in the striatum, SN, and PFC. We predicted aging would reveal the motor phenotype and accompany decreases in catecholamine levels, akin to human PD progression. Our findings revealed frank hyperkinetic behavior coincident with elevated DA in the SN and striatum in young adult PINK1 KO rats. With aging, but only nigral DA decreased by 18 months old in KO rats, coinciding with decreased locomotor activity that was restricted to the KO genotype. The hyperactive phenotype in young KO is speculated to arise from compensatory mechanisms responding to compromised PINK1 function.

## METHODS

### Animals

The **<u>first cohort</u>** consisted of male two-month-old Long-Evans Wild-type (WT, n=7) and PINK1 KO (n=7) rats aged in house to 16-months-old (Envigo, Madison, WI). The **<u>second cohort</u>** consisted of male young (6 to 7-month-old) WT (n=12; (Charles River)) and KO (n=14; (Envigo)) rats, and aged (18-month-old) WT (n=10; (n=5 from Envigo and n=5 from Charles River) and KO (n=10; (Envigo)) rats. All rats were singularly housed on a 12hr reverse light/dark cycle with food and water *ad libitum*. For one month prior to the experimental procedures, rats were handled at least three times a week and acclimated to their environment. We had two KO rats from our first cohort die unexpectedly, one at 7 and one 16 months while another drastically started losing weight at 10 months old. Additionally, we noted a sub-population of PINK1 KO rats experiencing hindlimb drag around the 7 to 8-month mark, which spontaneously resolved at the subsequent testing date. All experiments were performed after the protocol was approved by the Institutional Animal Care and Use Committee (IACUC) at the University of North Texas health Science Center and the Animal Care and Use Review Office (ACURO), Department of the Army, U.S. Army Medical Research and Development Council.

### Genotype confirmation

Genomic DNA was extracted from an ear hole punch from both KO and WT rats to confirm PINK1 gene knockout. The GenElute™ Mammalian Genomic DNA Miniprep kit was used with minor changes to the procedure. The full methodological procedure has been previously published by our lab (Salvatore et al., 2022)

### Locomotor Activity: Open-field test (OFT)

The open-field test (OFT) was used to capture and measure motor activity and was performed during the awake cycle using the Opto-Varimex-5 Auto-Track System (Columbus Instruments). Rats were each placed in a 17.5 x 17.5 x 12 in. acrylic chamber with clean bedding and allowed to freely explore the area under red light. The total distance traveled (in) and average speed (in/s) was captured for a period of one hour for three consecutive days. The median of the three days was used as the final score to calculate results. Rats from the **<u>first cohort</u>** of the study were tested monthly, starting at 3-months-old. Rats in the **<u>second cohort</u>** (young and aged) were tested for one session (median of 3 days) after handling procedures.

### Behavioral Test-Novel Object Recognition (NOR)

The Novel Object Recognition test assessed recognition memory and was conducted on the day following completion of the OFT. Rats were each placed alone in a 24” x 24”x 20” container under red-light and allowed to freely explore the area for 10 minutes. After this, the rats were removed, and two identical objects were placed in the corners equally spaced apart from the perimeters of the box. The rats were reintroduced into the testing area immediately and video recorded for 3 minutes. After 1 hour, one of the initial objects was placed back into the box along with a novel item of similar size but of different color and material. Rats were again placed into the box and video recorded for 3 minutes. Each rat was evaluated on the duration of time spent with the novel object. A discrimination index formula ((Novel-Familiar)/(Novel+Familiar)) was used to evaluate each rat’s ability to differentiate between the familiar and novel object. The **<u>first cohort</u>** was first evaluated monthly starting at 3-months with the initial set of objects, and then with a new set of objects starting at 12-months-old.

### Catecholamine and tyrosine hydroxylase assessment

All rats were euthanized within one week of last locomotor testing. The prefrontal cortex, striatum, ventral tegmental area (VTA), and substantia nigra (SN) were dissected on wet ice and immediately thereafter placed onto dry ice. Tissue was sonicated in perchloric acid/EDTA solution and spun to precipitate the protein pellet for further processing for quantitative blot immunolabeling of TH protein (Salvatore et al., 2012). The supernatant was then processed for analysis of dopamine (DA), norepinephrine (NE), and dihydroxyphenylacetic acid (DOPAC) using high-performance liquid chromatography (HPLC, for details on procedure, see Kasanga et al., 2023). The recovered protein pellet was further processed for western blot-based analysis and quantitation of TH protein (Millipore, Temecula, cat #: AB152; 1:1000 dil, CA, USA) against a calibrated TH protein standard to determine ng TH per µg total protein (Kasanga et al., 2023)

### Statistical Analysis

Statistical analysis was performed using GraphPad Prism (Version 10). A Student’s t-test was used to assess statistical differences between genotype at a single given time-point for the first cohort. A repeated measures two-way ANOVA was used to determine genotype and aging effects on motor results, matching month-to-month assessments, in the **<u>first cohort</u>**, with Bonferroni test used for post-hoc analysis. A two-way ANOVA was used to determine genotype and aging differences in the **<u>second cohort</u>**, with Bonferroni post-hoc analysis to correct for multiple comparisons. The Grubb’s test identified any outliers with alpha= 0.05, based on the *n* being evaluated.

## RESULTS

### Locomotor activity: Open-field test (OFT)

In our first cohort, PINK1 KO rats had greater total walking distance and average speeds from baseline (3 months) to 5 months old. However, this difference was not statistically significant. Starting at 6 months old, KO rats showed a significant decline, relative to their baseline scores, in these measures compared to WT rats (genotype: distance F(1,12)=10.63, *p=*0.0068; speed F=(1,12)=7.78, *p=*0.016) (age: distance F(12,129)=10.58, *p*<0.0001; speed F=(12,127)=2.92, p=0.0013) (age x genotype: distance F(12,129)=1.66, ns; speed F=(12,127)=1.88, p=0.042) **(Fig. 1A & B).** This decline was followed by a plateau and potentially an accelerated aging effect compared to WT rats. WT rats showed an expected initial decline from baseline beginning around 11 months old, consistent with the timing of aging-related hypokinesia reported in rat models of aging (Salvatore et al., 2016; 2017). In raw OFT scores, we identified a significant aging effect in both genotypes (distance F(13,144)=10.46, p<0.0001; speed F(13,146)=4.09, p<0.0001) and a significant age and genotype interaction (distance F(13,144)= 2.11, p=0.016; speed F(13,146)= 2.50, p= 0.004), yet we did not observe a genotype effect at any individual age (distance F(1,12)=0.006, ns; speed F(1,12)=0.36, ns) (**Fig. 1C & D**).

**Figure 1:**
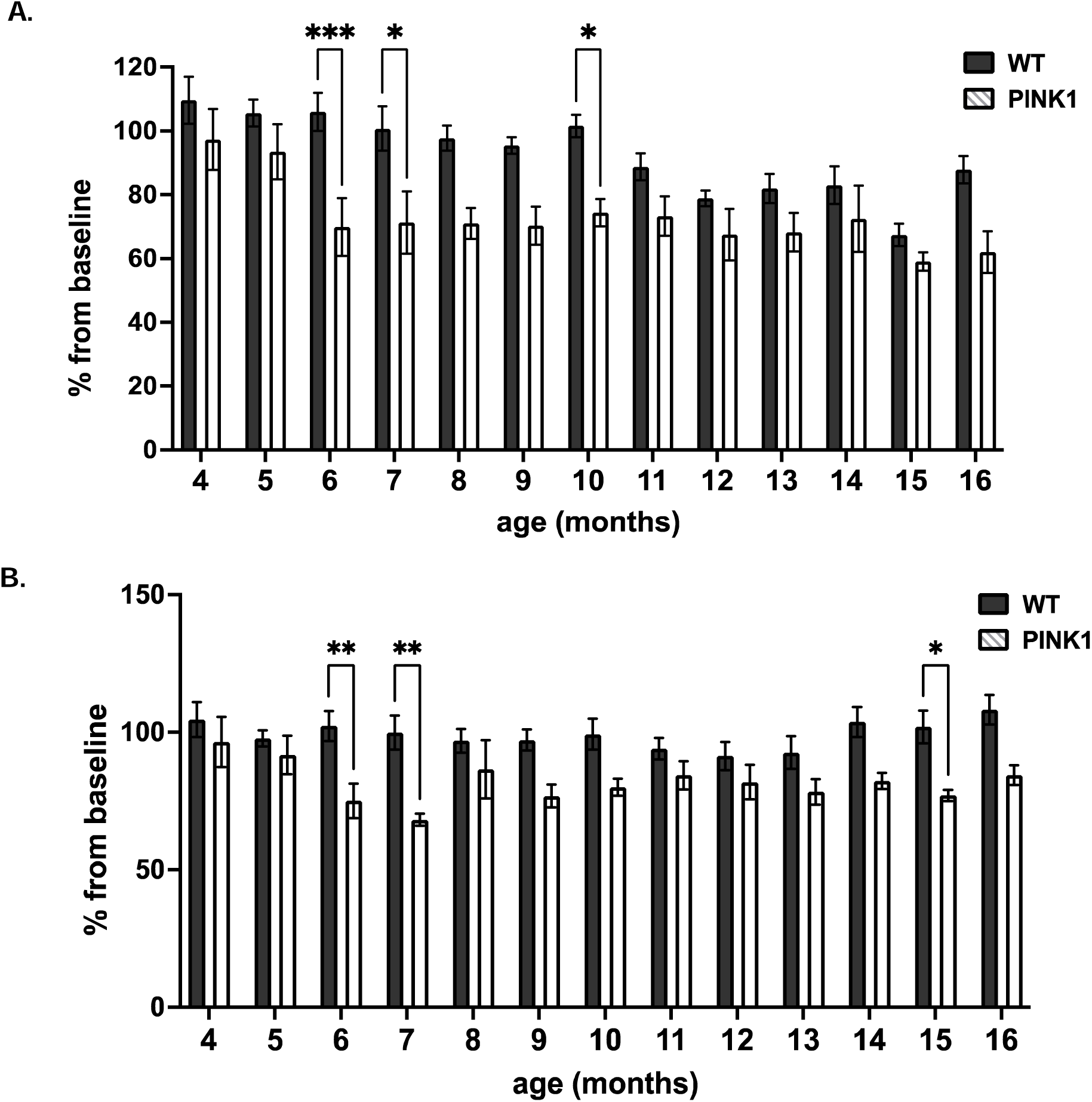
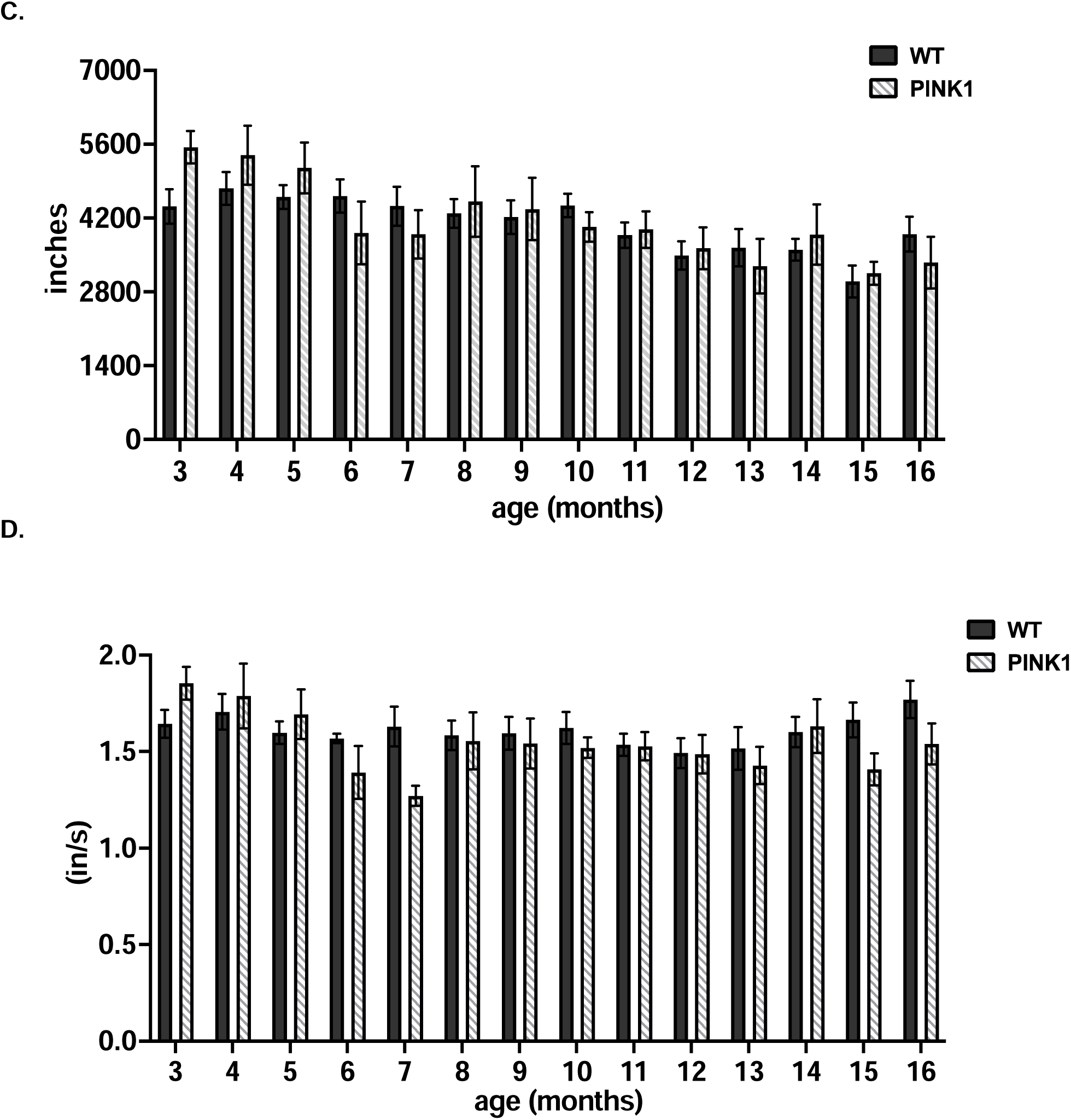
Open-field Test Results from First Cohort of Rats. In the first cohort, PINK1 KO rats showed a significant decrease in total distance traveled (DT) **(A)** and average speed **(B)**, compared to baseline results, starting at 6-months-old and continuing to 7 and then again at 11-months-old for DT and 7 and 15 months-old for speed. **(A)** DT: WT vs KO 4mo. (t= 1.40, ns, df= 141); 5mo. (t= 1.38, ns, df= 141); 6mo. (t= 4.12, *\*\*\*p*=0.0008, df= 141); 7mo. (t= 3.36, *\*p=* 0.01, df= 141); 8mo. (t= 2.72, ns, df= 141); 9mo. (t= 2.56, ns, df= 141); 10mo. (t= 3.31, \**p=* 0.01, df= 141); 11mo. (t= 1.99, ns, df= 141); 12mo. (t= 1.55, ns, df= 141); 13mo. (t= 1.63, ns, df= 141); 14mo. (t= 1.28, ns, df= 141); 15mo. (t= 1.05, ns, df= 141); 16mo. (t= 2.93, p=0.051, df= 141). **(B)** Speed: WT vs KO 4mo. (t= 1.03, ns, df= 139); 5mo. (t= 0.76, ns, df= 139); 6mo. (t= 3.47, *\*\*p*=0.009, df= 139); 7mo. (t= 3.56, *\*\*p=* 0.006, df= 139); 8mo. (t= 1.50, ns, df= 139); 9mo. (t= 2.26, ns, df= 139); 10mo. (t= 2.13, ns, df= 139); 11mo. (t= 1.41, ns, df= 139); 12mo. (t= 1.39, ns, df= 139); 13mo. (t= 1.98, ns, df= 139); 14mo. (t= 1.65, ns, df= 139); 15mo. (t= 3.06, *\*p=*0.03, df= 139); 16mo. (t= 2.93, p=0.051, df= 139). This decline from baseline was not seen in the WT control rats until 12-month-old. When evaluating raw scores though, there was no difference between PINK1 KO and WT rats in total DT **(C)** or average speed **(D)** at any tested age between 3 and 16 months old. **(C)** Raw DT: WT vs KO 3mo. (t= 2.12, ns, df=156); 4mo. (t= 1.91, ns, df= 156); 5mo. (t= 1.05, ns, df= 156); 6mo. (t= 1.32, ns, df= 156); 7mo. (t= 1.00, ns, df= 156); 8mo. (t= 0.29, ns, df= 156); 9mo. (t= 0.16, ns, df= 156); 10mo. (t= 0.87, ns, df= 156); 11mo. (t= 0.06, ns, df= 156); 12mo. (t= 0.13, ns, df= 156); 13mo. (t= 0.77, ns, df= 156); 14mo. (t= 0.52, ns, df= 156); 15mo. (t= 0.28, ns, df= 156); 16mo. (t= 0.96, ns, df= 156). **(B)** Raw Speed: WT vs KO 3mo. (t= 1.55 ns, df= 158); 4mo. (t= 0.61, ns, df= 158); 5mo. (t= 0.69, ns, df= 158); 6mo. (t= 1.55, ns, df= 158); 7mo. (t= 2.31, ns, df= 158); 8mo. (t= 0.21, ns, df= 158); 9mo. (t= 0.39, ns, df= 158); 10mo. (t= 0.76, ns, df= 158); 11mo. (t= 0.05, ns, df= 158); 12mo. (t= 0.11, ns, df= 158); 13mo. (t= 0.71, ns, df= 158); 14mo. (t= 0.30, ns, df= 158); 15mo. (t= 1.72, ns, df= 158); 16mo. (t= 1.53, ns, df= 158).

In our second cohort, young adult (7 month) KO rats exhibited significantly higher OFT scores in total walking distance (F(1,28)=10.07, p=0.0036) but not average speed compared to WT rats of the same age. However, in the 18-month-old group, we did not observe a significant difference in raw OFT scores between WT and KO rats. Aging-related decreases in total walking distance and average speed occurred only in KO rats between 7 and 18-months-old (distance F(1,28)=15.51, p=0.0005; speed (age x genotype interaction) F(1,28)=6.108, p=0.0198) **(Fig. 2A & B).**

**Figure 2:**
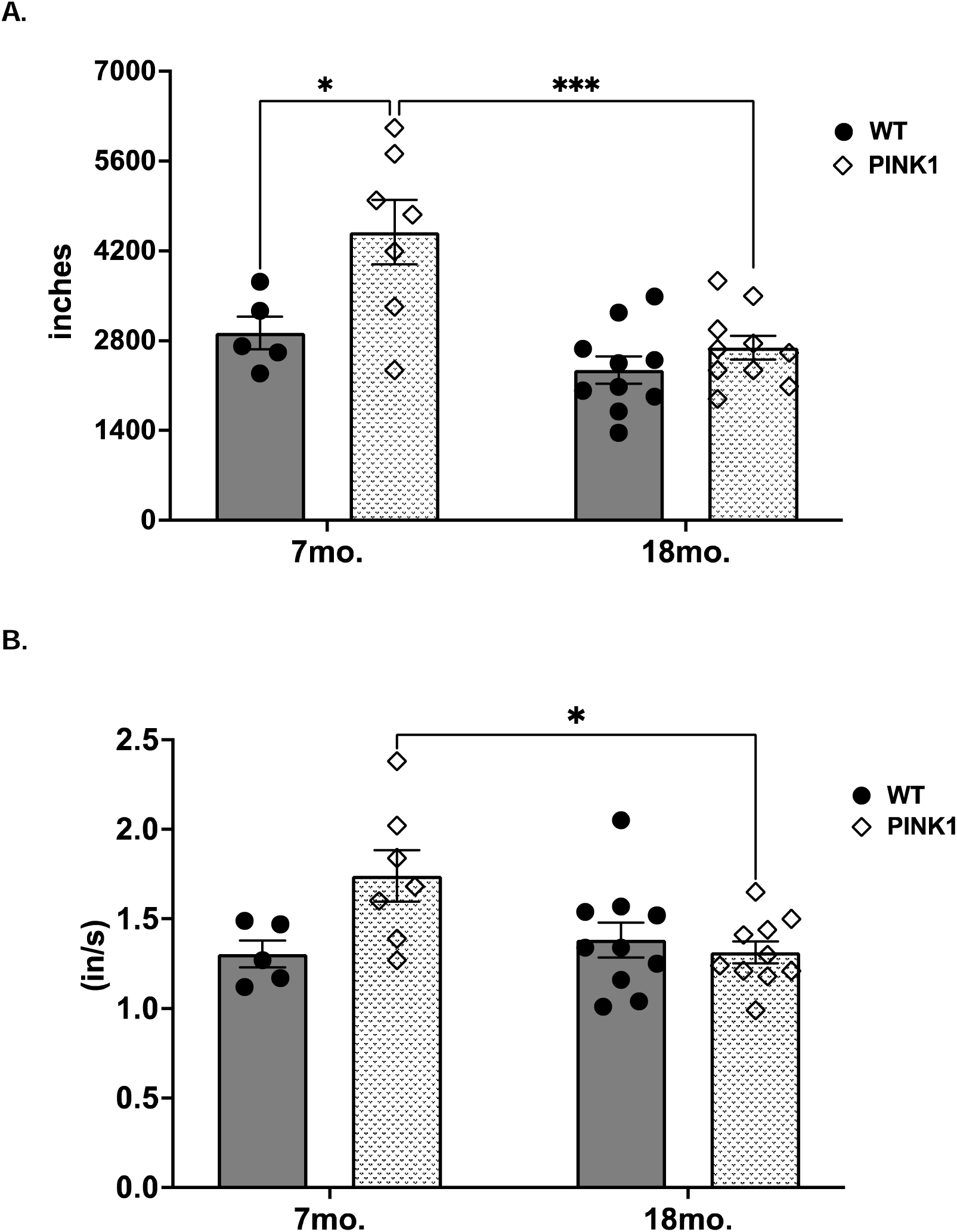
Open-field Test Results from Second Cohort of Rats. In the second cohort, young adult (7-month-old) PINK1 KO rats showed greater total distance traveled **(A)** than age-matched WT rats. This, along with average speed **(B)** significantly decreased by 18-months which did not occur between WT rats. Yet unexpectedly there was no difference in DT **(A)** or average speed **(B)** between 18-month-old WT and KO rats**. A. Distance:** 7mo. WT vs 7mo. KO (t=3.27, \**p=*0.017, df= 28); 7mo. WT vs 18mo. WT (t=1.28, ns, df= 28); 7mo. KO vs 18mo. KO (t=4.45, *\*\*\*p=*0.0007, df= 28); 18mo. WT vs 18mo. KO (t=0.95, ns, df= 28). **B**. **Speed**: 7mo. WT vs 7mo. KO (t=2.68, p=0.072, df= 28); 7mo. WT vs 18mo. WT (t=0.51, ns, df= 28); 7mo. KO vs 18mo. KO (t=3.12, \**p=*0.02, df= 28); 18mo. WT vs 18mo. KO (t=0.55, ns, df= 28).

### Recognition memory: NOR

In our first cohort there was no significant difference in discrimination index between KO and WT rats at 3 months old. Using the same set of objects, we did identify a significant age and genotype interaction (F(4,56)=4.296, p=0.004). Specifically, KO rats performing significantly worse than WT rats at 4 months old. However, this decline was not seen at subsequent testing times though **(Fig. 3A).** There was also no difference between genotypes when we tested the rats with a new set of objects at 12-months-old, and instead both KO and WT rats had a significantly higher discrimination score (F(1,22)=18.36, p=0.0003) from their initial 3 month NOR testing **(Fig. 3B).** However, we did observe overall higher levels of interaction with both familiar and novel objects in the KO rats, particularly starting at 12 months old **(Suppl. Fig. 2)**

**Figure 3:**
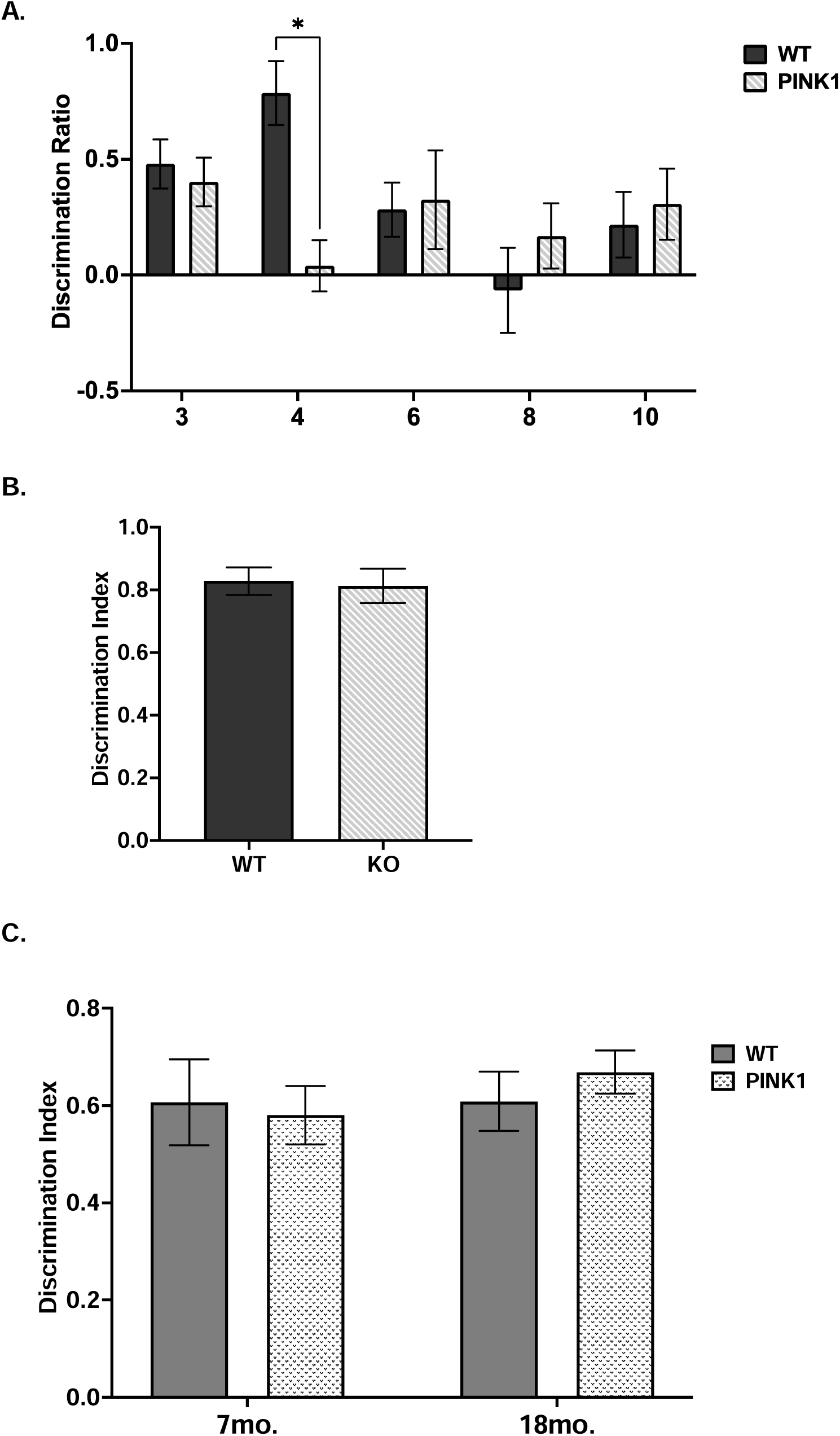
Novel Object Recognition Task. In the first cohort of rats **(A)**, KO rats exhibited a significantly lower discrimination score at 4 mo. compared to WT rats, a difference not found at baseline or in subsequent testing. WT vs KO 3mo. (t= 0.53, ns, df= 12); 4mo. (t= 4.23, *\*p*=0.012, df= 12); 6mo. (t= 0.17, ns, df= 12); 8mo. (t= 1.01, ns, df= 12); 10mo. (t= 0.42, ns, df= 12). There was also no difference between genotypes when the rats were assessed on a new set of objects **(B)** at 12mo. (t= 0.21, ns, df= 10). In the second cohort **(C),** there was no significant genotype or aging effect between 7-month-old and 18-month-old PINK1 KO and WT rats as far as ability to discriminate between a familiar and novel object. 7mo. WT vs 7mo. KO (t= 0.28, ns, df= 27); 7mo. WT vs 18mo. WT (t=0.02, ns, df= 27); 7mo. KO vs 18mo. KO (t= 1.07, ns, df= 27); 18mo. WT vs 18mo. KO (t= 0.75, ns, df= 27).

In our second cohort, we used the same set of objects as in the first cohort and likewise we did not find a significant difference in discrimination index between genotypes in either the 7-month or 18-month-old groups. We also found that both WT and KO 18-month-old rats had a significantly higher discrimination score compared to their younger counterpart which likely pertains to task object preference versus a pathological indication **(Fig. 3B).**

### DA Tissue Content and DA Turnover Rate

At 16 months of age, the longitudinal (first) cohort of PINK1 KO rats did not show any significant difference in DA content compared to age-matched WT in the SN (t=0.60, ns, df=11**) (Fig. 4A),** striatum (t=1.67, ns, df=10) **(Fig. 4B),** or VTA (t=0.08, ns, df=10) **(Suppl. Fig.1A)**. However, in the PFC, DA tissue levels were greater in KO compared to WT rats (t=2.97, p=0.01, df=10) **(Fig. 4C)**. We did not find significant differences in DA turnover between KO and WT in the SN (t=0.35, ns, df=11) or striatum (t=1.52, ns, df=11) (**Fig. 5A & B**).

**Figure 4:**
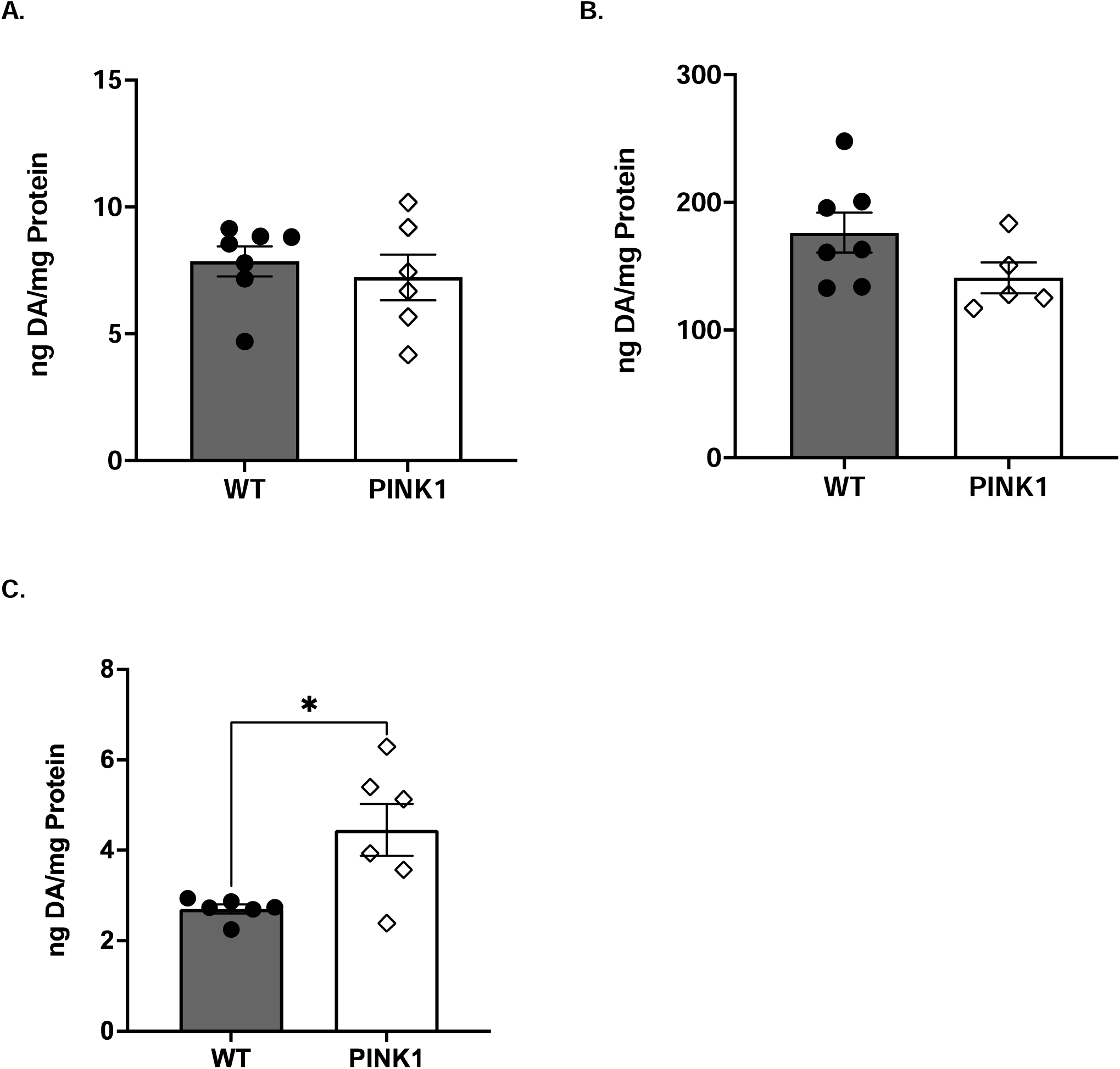
DA Tissue Content First Cohort. In the first cohort there was no significant difference in dopamine (DA) content in the substantia nigra (SN) **(A)** (t=0.60, ns, df=11) or in the striatum **(B)** (t=1.67, ns, df=10) of 16-month-old WT compared to KO rats. Yet, there was significantly higher DA level in the prefrontal cortex (PFC) **(C)** of 16-month-old KO rats compared to age-matched WT rats (t=2.97, \**p*= 0.01, df=11)

**Figure 5:**
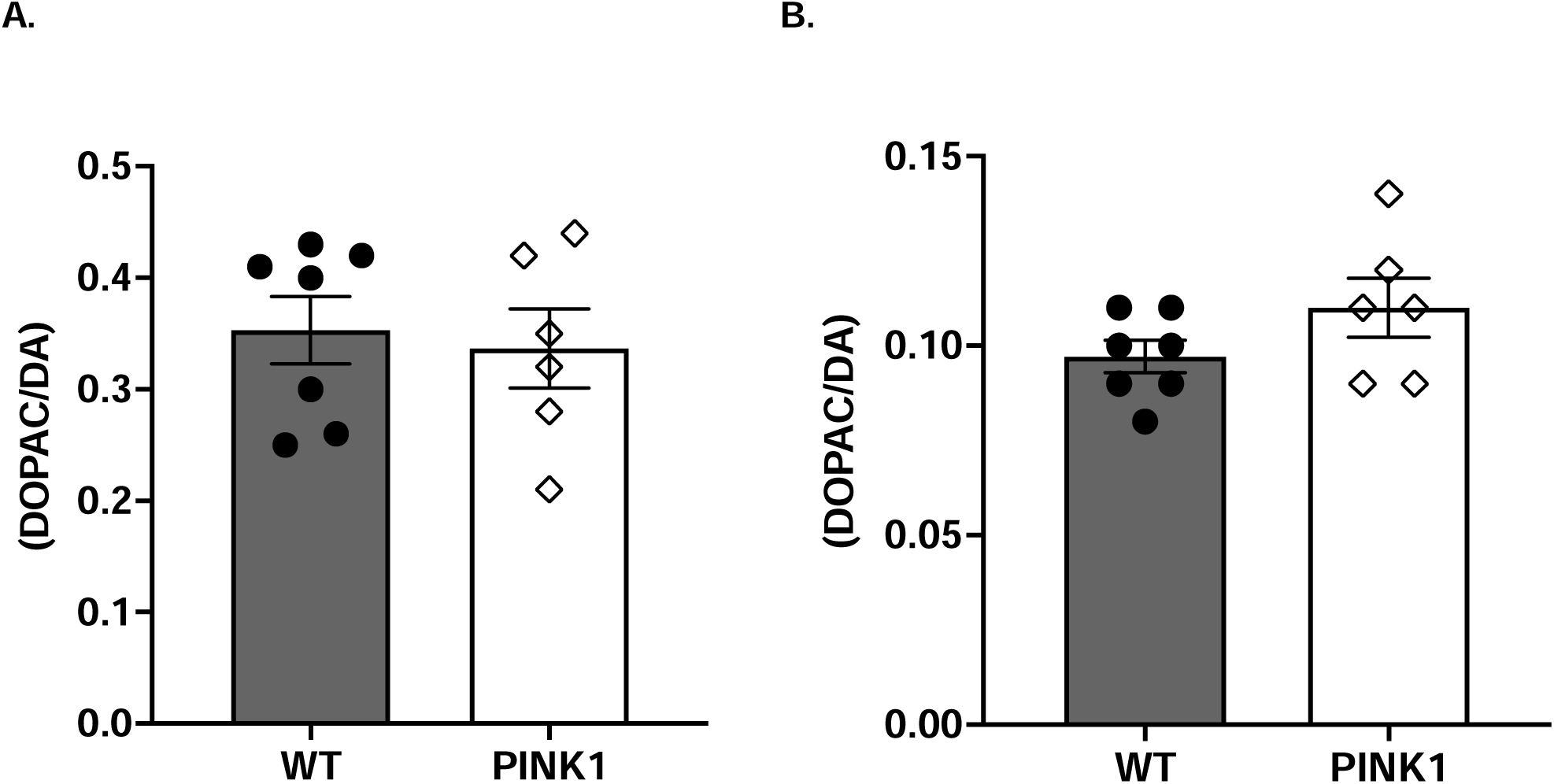
DA Turnover Rate First Cohort. In the cross-sectional study there was no differences between genotypes in DA turnover rate (DA/DOPAC) in either the substantia nigra **(A)** (t= 0.35, ns, df= 11) or in the striatum **(B)** (t= 1.52, ns, df= 11).

In the cross-sectional (second) cohort, there was a significant genotype effect (F(1,29)=8.80, p=0.006) and interaction of age and genotype in DA levels in the SN (F(1,29)=9.11, p=0.005). This difference was due to greater DA content in 7-month-old PINK1 KO rats compared to age-matched WT controls **(Fig. 6A)** and an aging-related decrease that was restricted to the KO. Notably, however, there was no significant difference in nigral DA content between 18-month-old KO and WT rats, similar to the results from the longitudinal first cohort.

**Figure 6:**
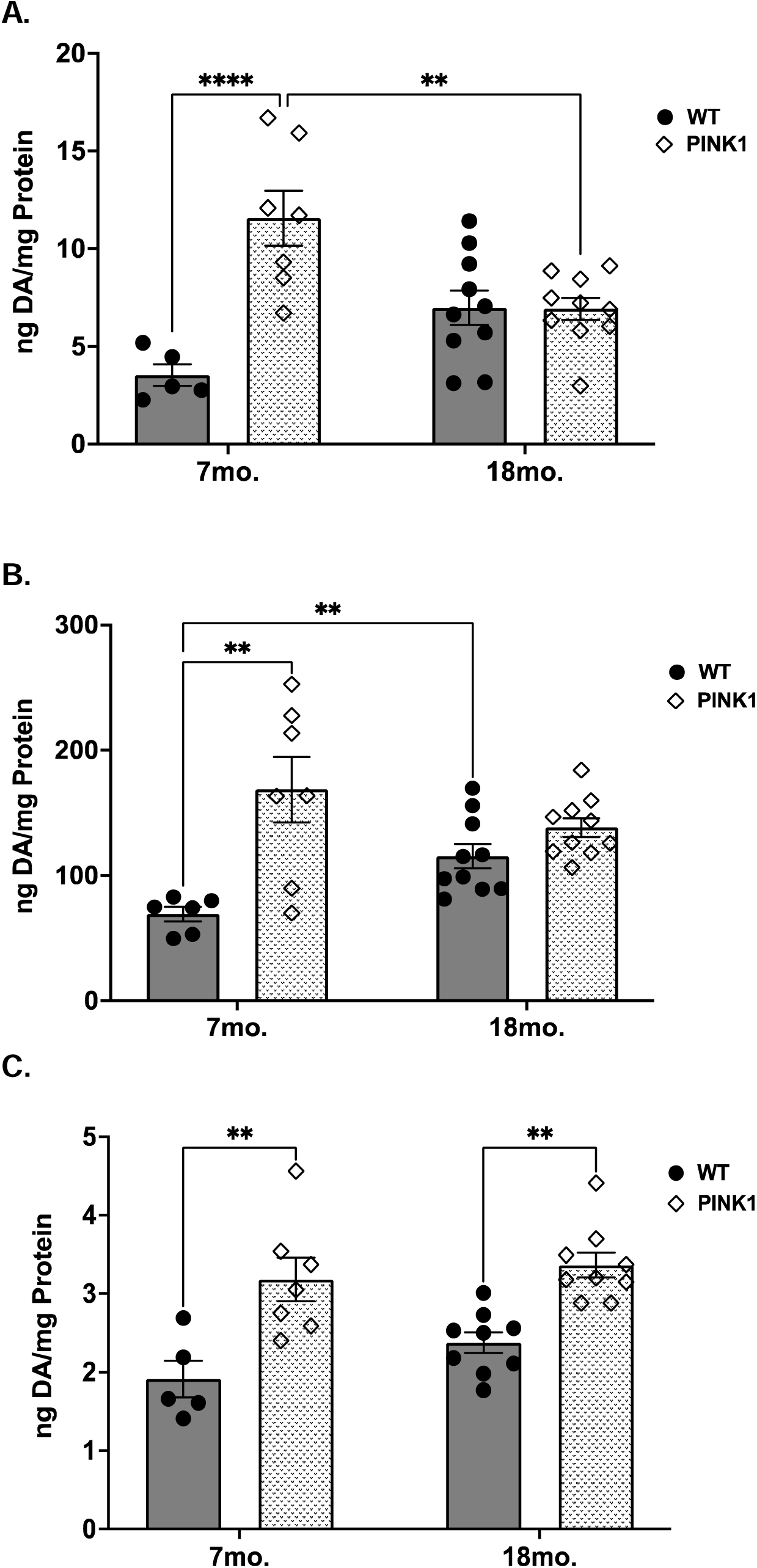
DA Tissue Content Second Cohort. In the second cohort, there was significantly higher DA in the SN **(A)** and the striatum **(B)** of 7-month-old KO rats compared to WT controls, with no genotype difference in either region between 18-month-old rats. There was also a significant decrease in 18-month-old rats in the SN but not the striatum in which instead there was a significant increase in DA between 7-mointh-old and 18-month-old WT rats. **(A)** SN: 7mo. WT vs 7mo. KO (t= 5.27, *\*\*\*\*p*< 0.0001, df= 28); 7mo. WT vs 18mo. WT (t= 2.42, ns, df= 28); 7mo. KO vs 18mo. KO (t= 3.62, **p= 0.0069, df= 28); 18mo. WT vs 18mo. KO (t= 0.05, ns, df= 2). **(B)** Striatum: 7mo. WT vs 7mo. KO (t= 3.85, *\*\*p*= 0.0035, df= 29); 7mo. WT vs 18mo. WT (t= 3.94, **p= 0.0028, df= 29); 7mo. KO vs 18mo. KO (t= 1.54, ns, df= 29); 18mo. WT vs 18mo. KO (t= 1.95, ns, df= 29). Similarly, to the first cohort, there was significantly higher DA in the PFC **(C)** between both 7-month-old and 18-month-old KO rats compared to their age-matched WT controls, yet there was no aging effect. 7mo. WT vs 7mo. KO (t= 4.07, *\*\*p=* 0.0023, df= 26); 7mo. WT vs 18mo. WT (t= 1.56, ns, df= 26); 7mo. KO vs 18mo. KO (t= 0.68, ns, df= 26); 18mo. WT vs 18mo. KO (t= 3.94, **p= 0.0033, df= 26).

In the striatum, there was a significant genotype effect (F(1,29)= 17.87, p=0.0002) with higher DA in 7-month-old PINK1 KO compared to same age WT and an aging effect (F(1,29)=15.39, *p=*0.0005), with increased levels from young to aged that was restricted to the WT **(Fig. 6B).**

We found no significant age (F(1,28)=0.52, ns) or genotype effect (F(1,28)=0.88, ns) in the VTA **(Suppl. Fig.1B).** Similar to the results from our first cohort, we identified a significant genotype effect on DA content in the PFC (F(1,26)=31.86, p<0.0001) where both young and aged PINK1 KO rats exhibited significantly higher DA content than their age-matched WT counterparts, although there was no effect of aging in either genotype **(Fig. 6C).** Moreover, we did not find any significant differences in DA turnover between genotypes in the SN **(Fig. 7A),** but did find a significant interaction (F(1,29)=11.93, *p=*0.0017), aging (F(1,29)=24.69, *p<*0.0001), and genotype effect (F(1,29)=13.90), *p=*0.0008) in the striatum which was due to an increase between 7-month-old WT to KO rats and reduction in DA turnover between 7 and 18-month-old WT rats **(Fig. 7B)**.

**Figure 7:**
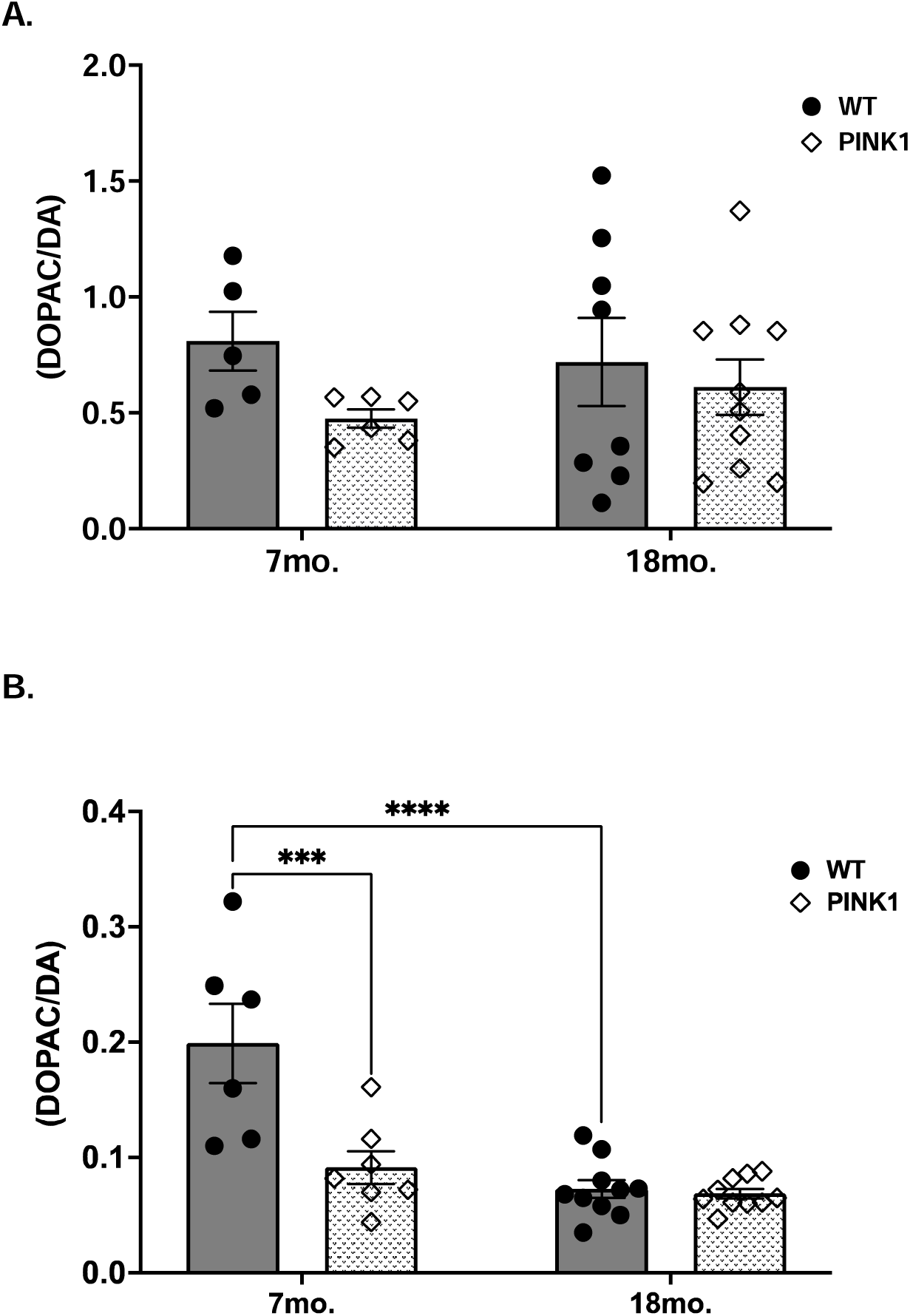
DA Turnover Rate Second Cohort. For DA turnover rate in this second cohort, there was no genotype or age difference in the SN **(A)** but there was a significant higher rate in 7-month-old WT rats compared to KO and decrease in the striatum **(B)** only between 7-month-old and 18-month-old WT rats. **(A)** SN: 7mo. WT vs 7mo. KO (t= 1.43, ns, df= 25); 7mo. WT vs 18mo. WT (t= 0.41, ns, df= 25); 7mo. KO vs 18mo. KO (t= 0.68, ns, df= 25); 18mo. WT vs 18mo. KO (t= 0.59, ns, df= 25). **(B)** Striatum: 7mo. WT vs 7mo. KO (t= 4.61, ***p= 0.0005, df= 29); 7mo. WT vs 18mo. WT (t= 5.82, *\*\*\*\*p*< 0.0001, df= 29); 7mo. KO vs 18mo. KO (t= 1.09, ns, df= 29); 18mo. WT vs 18mo. KO (t= 0.22, ns, df= 29).

### NE Tissue Content in the PFC

In the first cohort, there was a significant decrease in NE content in the PFC of 16- month-old KO rats compared to WT rats (t=2.31, *p=*0.046, df=9) **(Fig. 8A).** On the other hand, in our second cohort an aging-related effect was not detected (F(1,26)=0.42, ns), but a significant genotype effect was indicated (F(1,26)=10.05, p=0.0039) which was due to a higher NE content in the 7-month-old KO rats compared to the age-matched WT rats. We also identified a significant interaction of age and genotype (F(1,26)=7.50, p=0.011) **(Fig. 8B).**

**Figure 8:**
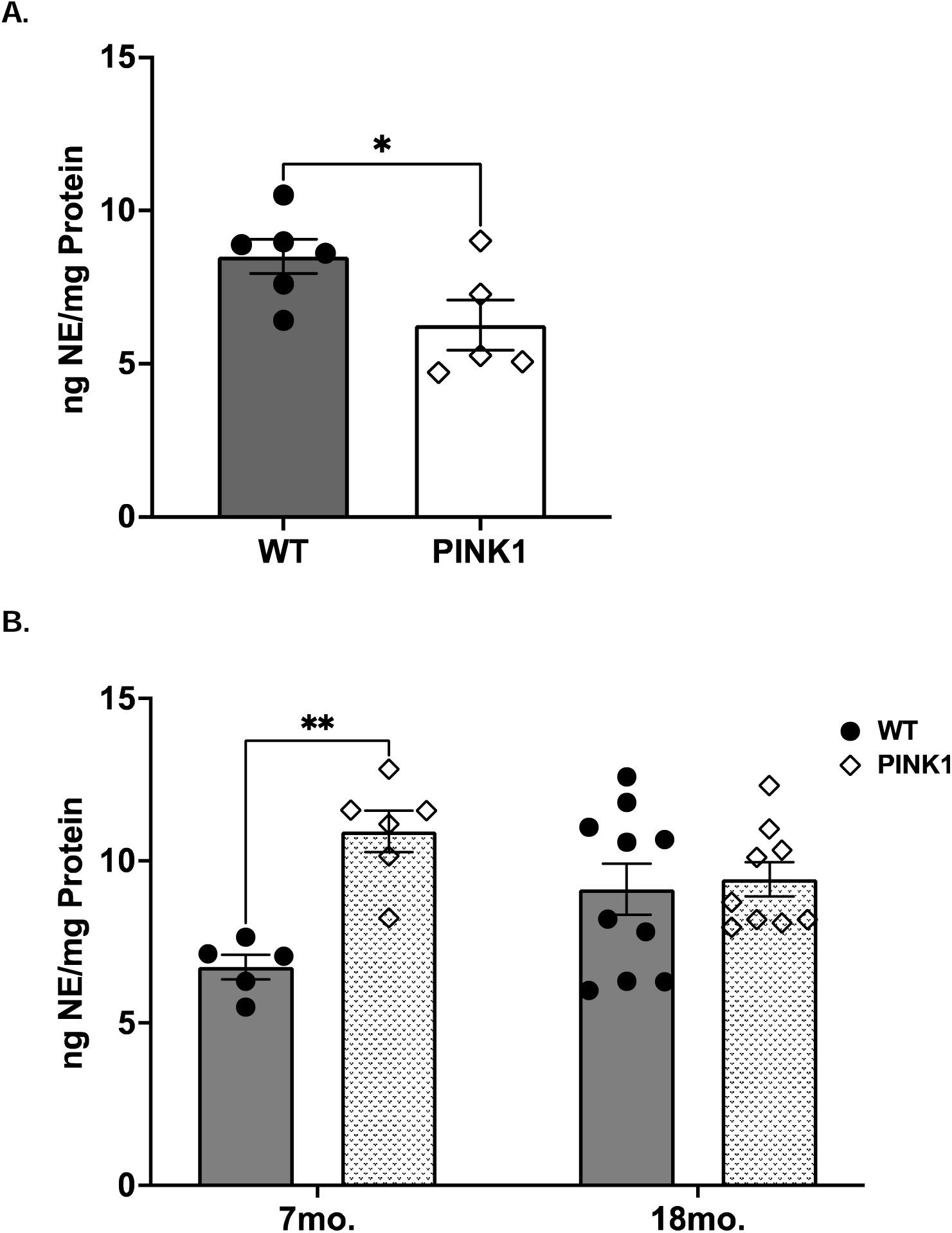
NE in the PFC. In the first cohort of rats, there was about a 20% reduction of NE in the PFC **(A)** of 16-month-old KO rats compared to age-matched WT rats (t= 2.31, \**p*= 0.046, df= 9). In the second cohort, there was significantly higher NE in the PFC **(B)** of 7-month-old KO rats compared to age-matched WT controls, but there was no age effect or genotype difference between 18-month-old rats. 7mo. WT vs 7mo. KO (t= 3.71, **p= 0.006, df= 26); 7mo. WT vs 18mo. WT (t= 2.35, ns, df= 26); 7mo. KO vs 18mo. KO (t= 1.51, ns, df= 26); 18mo. WT vs 18mo. KO (t= 0.35, ns, df= 26).

### Tyrosine hydroxylase (TH) Protein Expression

In the longitudinal study (first cohort), there was no significant difference in TH protein levels in either the SN (t=0.94, ns, df=10) or the striatum ((t=0.80, ns, df=9) at 16 months old **(Fig. 9A & B)).**

**Figure 9:**
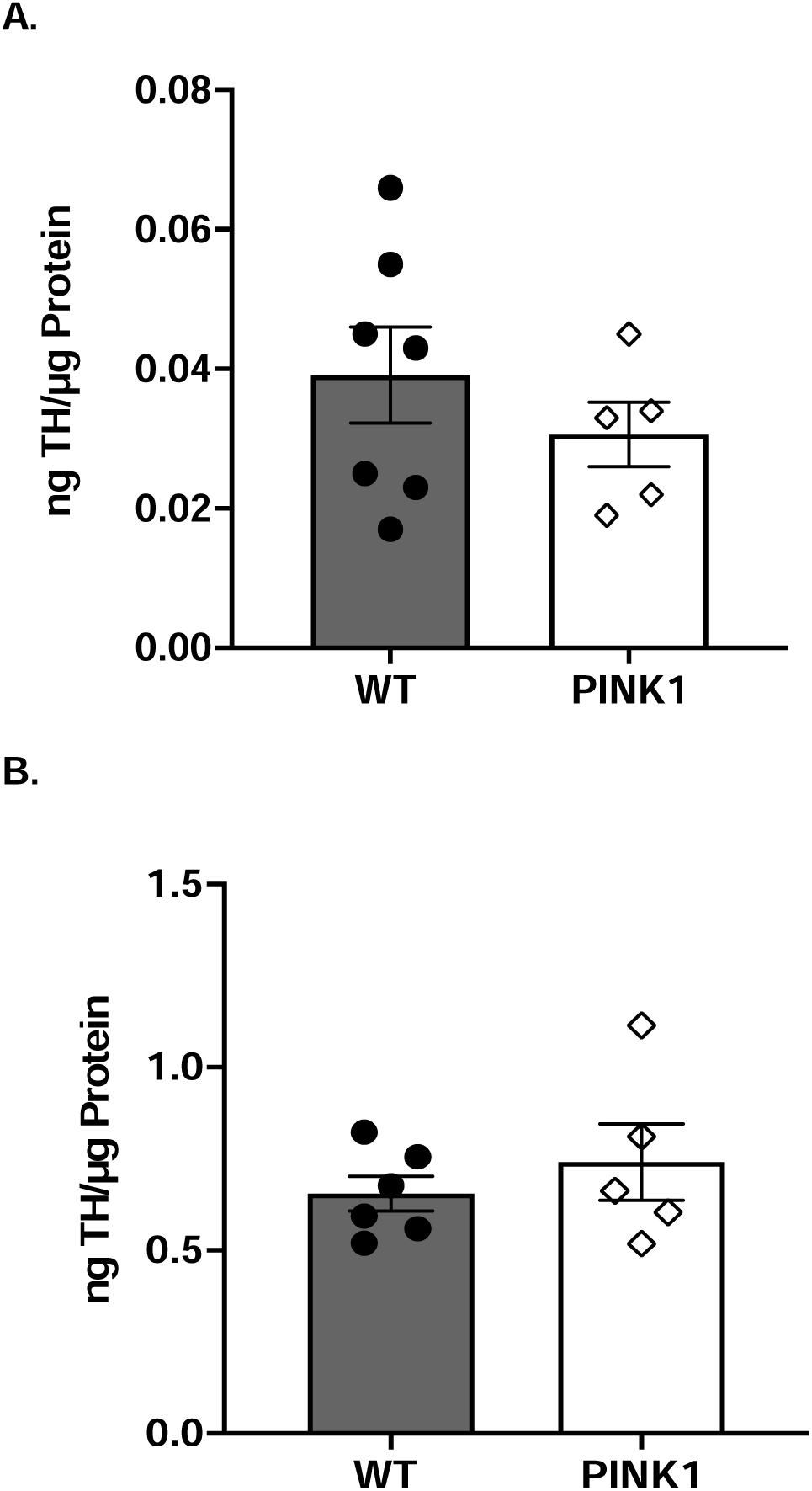
TH expression in SN and Striatum in First Cohort. While there was a a slight decrease in total tyrosine hydroxylase (TH) in the SN **(A)** of the first cohort this was not significant (t= 0.94, ns, df= 10). There was also no change in the striatum **(B)** of 16-month-old WT versus PINK1 KO rats (t= 0.84, ns, df= 9).

In the cross-sectional study (second cohort), there was a significant aging effect (F(1,26)=16.09, p=0.0005) and interaction between age and genotype (F(1,26)=5.718, p=0.024) in the SN which was due to decreased TH protein expression between 7 and 18-month-old KO rats **(Fig. 10A).** We also found a significant interaction of age and genotype in striatal TH levels (F=(1,29)=6.881, p=0.0137), but no differences in TH between 7-month or 18-month groups **(Fig. 10B).**

**Figure 10:**
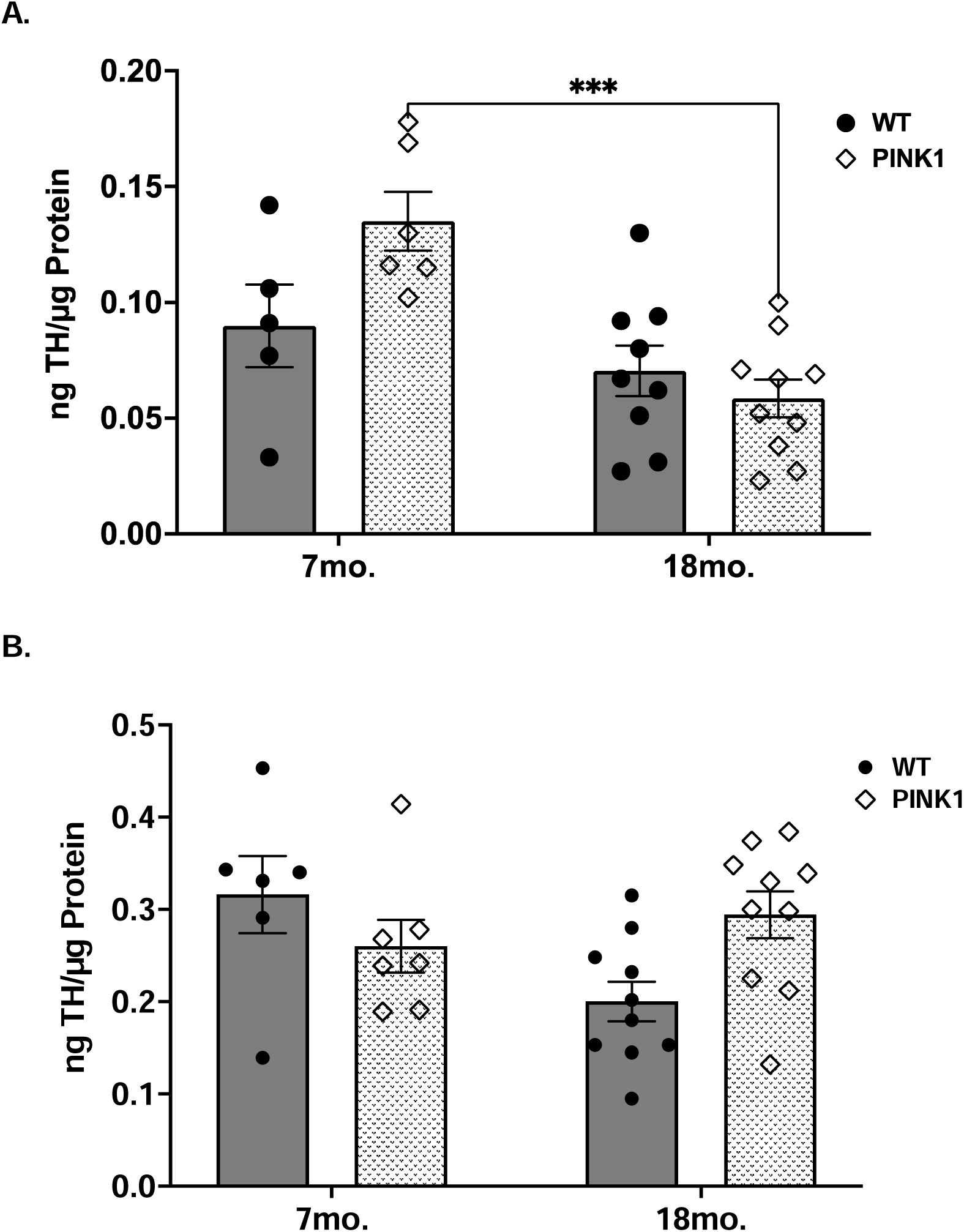
TH expression in SN and Striatum in Second Cohort. In the cross-sectional (second) cohort, there was a significant decrease in TH protein the SN **(A)** between 7-month-old and 18-month-old rats KO rats, which was a decline not found in the WT rats. 7mo. WT vs 7mo. KO (t= 2.37, ns, df= 26); 7mo. WT vs 18mo. WT (t= 1.10, ns, df= 26); 7mo. KO vs 18mo. KO (t= 4.71, *\*\*\*p*= 0.0003, df= 26); 18mo. WT vs 18mo. KO (t= 0.83, ns, df= 26). There was no age or genotype effect in the striatum **(B**), yet there was a significant age x genotype interaction. 7mo. WT vs 7mo. KO (t= 1.28, ns, df= 29); 7mo. WT vs 18mo. WT (t= 2.81, ns, df= 29); 7mo. KO vs 18mo. KO (t= 0.85, ns, df= 29); 18mo. WT vs 18mo. KO (t= 2.59, ns, df= 29).

## DISCUSSION

Aging is the number one risk factor for PD (Collier et al., 2011; 2017). Although familial forms of PD typically manifest at an earlier age than idiopathic PD (Hayashida et al., 2021), aging still influences the timing of phenotypic expression. For a PINK1 mutation, the median age of disease onset in homozygote or mutation carriers is estimated to be 30 to 39 years old (Valente et al., 2004; Bonifati et al., 2005; Ishihara-Paul et al., 2008; Hayashida et al., 2021). Therefore, it stands to reason that the lack of consistency in presentation of a motor phenotype in the PINK1 KO rat model may be that most studies are conducted primarily between 4 and 8 months old (Grant et al., 2015; Ferris et al., 2018; Cai et al., 2019; de Haas et al., 2019; Creed et al., 2019; Grigoruţă et al., 2020). In reference to human age, ages 4 to 8 months old correlate with teen to mid-20s; which is under the age of presentation of ∼35 in most humans (Quinn, 2005). We found evidence of a highly significant influence of aging when comparing Pink1 KO against the respective age-matched WT control in both a longitudinal and cross-sectional study.

Only one study to our knowledge has thus far conducted assessment of DA signaling beyond 8 months old, and in that study a moderate decrease in striatal DA release was reported in 14 month old Pink1 KO mice (Zhi et al., 2019). Accordingly, our study represents a rigorous longitudinal and cross-sectional investigation of aging-related timing of motor and cognitive impairments against the catecholamine profile in striatum and three additional DA regions at advanced age (beyond 12 months). By integrating aging into the study, our findings contribute to refining the translatability of this model in PD pre-clinical research. This is particularly important given the inconsistent results from previous research on this genetic model.

Thus, our locomotor results contradict with some prior reports (Dave et al., 2014; de Haas, 2019), and also reveal a high degree of heterogeneity in the timing of motor decline onset. Contrary to expectations, biological aging coupled with the PINK1 mutation did not produce parkinsonian-like bradykinesia, in terms of raw scores, as compared to age-matched WT rats. That is, the basis for a significant decrease in locomotor activity attributed to aging in the PINK1 KO rats was driven solely by hyperactivity at a young age. Hyperactivity in the young PINK1 KO rats has been previously reported in one additional study (Grigoruţă et al., 2020), which noted the appearance of a bradykinetic phenotype only under psychological stress. In all, we did not observe a significant decrease in raw motor activity in the KO when compared to WT rats at any specific age. While this was the case using the open-field test, which measures spontaneous activity, others measuring balance and grip strength have identified significant differences (Grant et al., 2015; Ferris et al., 2018; Dave et al., 2014). While this could suggest that PINK1 KO rats represent a decline in fine motor skills associated with PD, we expected to see gross locomotor differences, particularly bradykinesia or hypokinesis with advanced age.

Overall, the motor findings in this study, along with our previous results using this model (Salvatore et al., 2022) indicate that young adult PINK1 KO rats possibly undergo a compensatory phase of heightened activity before evident motor decline (Kasanga et al., 2023). This hyperactivity of young Pink1 KO rats has only been reported twice in previous studies (Ferris et al., 2018; Salvatore et al., 2022), but has yet to be reported in human PD, suggesting that the rat species may respond differently to a PINK1 mutation than humans. Given the sheer number of Pink1 variants in human, most frequently involving point or frameshift mutations, it may prove difficult to capture a translational phenotype (Kasten et al., 2018; Ma et al; MDS Gene Database). This model has a 26-basepain frameshift mutation, presumably creating a premature stop codon (Dave et al., 2014). However, the extent of this mutation may not be sufficient to drive intended nigrostriatal loss (Klein et al., 2007) without additional risk factors of PD such as lifestyle or environmental stressors and exposures. Although aging is the number 1 risk factor for PD, our results show despite strong interaction of aging with genotype, the overt motor phenotype of bradykinesia is simply not present.

While PD is a multifactorial disease, with pathological changes including neuroinflammation, increased oxidative stress, and ubiquitin-proteasome system failure, the defining and distinctive pathology that leads to the disease is a loss of dopaminergic neurons in the SN (McDonald et al., 2018; Hartmann et al., 2004; McNaught et al., 2001; Hirsch et al., 2003). Regardless of whether PD is idiopathic or of familial origin, postmortem studies show a variable 50-90% DA cell loss in the SN, while >80% loss in the striatum is associated with the onset of characteristic motor phenotype (Kordower et al., 2013; Nybo et al., 2020; Mazumder et al., 2022; Heng et al., 2023). Arguably, rodent PD models must reflect the human nigrostriatal pathologies. The cross-sectional study did reveal a significant decrease in DA and total TH levels in the SN of PINK1 KO rats from 7 to 18 months, a decrease that was not present in WT controls. This decline in DA was specific to the SN as we did not find similar changes in the striatum or VTA. In terms of an aging effect, we did not find a significant difference in DA or TH in the SN when strictly comparing 16-month or 18-month-old KO rats to age-matched WT controls. This result was found in addition to no decreases in striatal TH or DA, which has been reported in most PINK1 KO studies (Dave et al., 2014; Grant et al., 2015; Villeneuve et al., 2016; Creed et al., 2019). Thus, at the very least, our rigorous evaluation of this KO model suggests that it falls short of emulating the characteristics of nigrostriatal neuron loss, DA regulation, and motor phenotype seen in human PD.

Nonetheless, it is especially important to note the congruity of changes in nigral DA levels against the hyperactivity in young Pink1 KO rats versus the young WT, in addition to the aging-related changes in the SN that were restricted to the KO with aging. The hyperactivity was associated with increased DA in the KO versus WT in SN and striatum. However, the aging-related decrease in locomotor activity in the KO was coincident with decreased nigral, but not striatal, DA and TH protein. The overwhelming data gathered from our group and one other study show evidence that nigral DA signaling does influence specific locomotor functions related to movement initiation (Pruett and Salvatore, 2013; Salvatore et al., 2019; 2023; González-Rodríguez et al., 2021; Kasanga et al., 2023). Moreover, the aging-related change in the SN of the KO was not seen in the striatum, consistent with discrepancies reported in aging and PD models alike (Salvatore et al., 2024). The unexpected increase in striatal DA in WT rats from 7 to 18-months may be related to the source of Long-Evans rats, which were sourced from Charles River and Envigo. Studies have indeed reported animal sourcing may affect biological outcomes (Oliff et al.; 1995; Khoo et al., 2022). Nonetheless, KO and WT differences were not found in the aged groups, regardless of vendor. Therefore, the lack of SN and striatal DA or TH loss in this model, when compared against WT levels, is inconsistent with human PD, as is the lack of difference in raw locomotor activity between WT and KO at older age. These findings do not show fidelity to the conditions of slow progression of motor phenotype and nigrostriatal pathology reflecting human PD. Unfortunately, this rigorous evaluation leaves us to conclude that there are significant limitations of this model at least in terms of translation to human PD: and thus the need to develop a model that captures both the nigrostriatal neuron loss, impaired DA regulation, and slower rate of motor decline continues.

Apart from movement related symptoms, about 30% of PD patients will develop mild cognitive impairment which may present as a subtle decline in executive functioning (Aarsland et al., 2017; Durcan et al., 2019; Nejtek et al., 2021). We propose that replicating the non-motor symptoms of PD in animal models can improve translation and can lead to mechanistic insights to advance treatment options in the earliest stages of the disease. Addressing PD at its earliest stages is essential, as diagnostic indices in the prodromal stage have not been standardized and no pharmacological treatment options are currently available to address these symptoms. We assessed cognitive changes in this model using the NOR task, which uses a rat’s innate curiosity to measure recognition memory, to identify if cognitive decline occurs alongside with progression of the disease as could be expected in human PD (Ennaceur et al., 2010). We only found a significant difference in discrimination between a familiar and novel object between the genotypes at 4 months old, but not at 3, 7, 12, or 18 months. This partially aligns with recent findings by Pinizzotto et al. (2022) who did not identify recognition memory deficits in PINK1 KO male rats at 3 months but did at 5, 7, and 9 months.

In line with Pinizzotto (2022), we observed that, on average, PINK1 KO rats spent more total time, particularly at older ages, interacting with objects compared to WT rats. While this observation is not a scored parameter of the task, it gives insight into some evident behavioral differences within this model. While this increased interactivity could suggest decreased anxiety, it could be driven by DA dysregulation, evidenced by increased DA in the PFC, leading to decreased habituation or perseveration (Sun et al., 2013; Murphy et al., 1996: Ren et al., 2021; Salvatore et al., 2021). Yet, two studies investigating cognitive functioning in this model have failed to note any significant deficits (Cai et al., 2019; Ferris et al., 2018). Nonetheless, the similar NOR scores may be attributed, at least in the younger cohorts, to the task’s inability to capture subtle changes in executive functioning, the domain of cognition initially impacted in PD (Ferris et al., 2018; Dirnberger et al., 2013; Aarsland et al., 2017; Nejtek et al., 2021; Soto et al., 2024). F-FDOPA PET scans have suggested that elevated dopaminergic activity in the PFC and constitutes a pathological feature of early-stage PD (Bruck et al., 2005; Kaasinen et al., 2001; Fallon et al., 2012), which is in alignment with our DA results in PFC. Taken together, the Pink1 KO result suggests early compensatory mechanisms in SN and PFC are present in this model, evidenced by DA and NE dysregulation in these regions.

Most studies with Pink 1 KO rat model indicate moderate loss of DA neurons, although the precise age of onset, degree of loss, and progression rate of decline varies (Dave et al., 2014; Villeneuve et al.2016; de Haas et al., 2019, Grigoruta et al., 2020, DeAngelo et al., 2022). This inconsistency across studies may be attributed, in part, to differences in methodologies, environmental factors, or spontaneous genetic mutations within the model. Nonetheless, there is evidence of specific non-motor and sensorimotor features that are reminiscent of prodromal or early-stage PD, including vocalization and learning deficits, and alpha-synuclein and mitochondrial dysregulation (Maynard, 2020; Pinizzottoo et. al, 2022; Kelm-Nelson et al., 2021; Creed et. al, 2019; Villeneuve et al., 2016; Cai et al., 2019; Hoffmeister et al., 2020; Grant et al., 2015; Cullen et al., 2018; Lechner et al., 2022; Soto et al., 2024). Accordingly, this highlights the apparent complexities related to PINK1 function and its influence on nigrostriatal neurodegeneration and DA regulation (Urrutia, 2014). Therefore, as with most PD models, the PINK1 KO rat may prove useful for evaluating certain early symptoms and pathological changes of PD. Our results underscore the need for studies to account for the existing heterogeneity and limitations in the PINK1 KO model and consider potential compensatory mechanisms during the early stages of its lifespan that counteract the influence of the mutation, and thus possibly mask a parkinsonian phenotype. That said, we did expect aging to accelerate and reveal the parkinsonian phenotype as compared to wild-type, which did exhibit

### Conclusion

This study provides a rigorous evaluation of the influence of aging on behavioral and related catecholamine-driven mechanisms in PINK1 KO rats and highlights the challenges in using this PD model. Inconsistencies in behavioral and neurobiological characteristics have raised questions about its suitability as a preclinical model. We aimed to bridge this gap by incorporating the role of aging in motor decline and its relevance to the progressive nature of PD. However, our findings reveal a high degree of variability in the timing of locomotor motor impairment onset in the lifespan. Contrary to expectations, the combination of biological aging and the PINK1 mutation did not align with the expected aging-related impairments observed in human PD. Instead, our results suggest a potential compensatory response of heightened activity and increased DA in SN of young adult against the influence of PINK1 mutation. Furthermore, we did not find a consistent decline in motor activity, and cognitive assessments did not reveal sustained differences between young or aged PINK1 KO and WT rats. These inconsistencies, combined with variations in the loss of dopaminergic neurons and the complexity of PINK1 mutations in humans, underscore the limitations of this model for replicating the slow progression of motor symptoms and nigrostriatal pathology distinctive of human PD. Thus, it is crucial to acknowledge the heterogeneity and constraints of the PINK1 KO rat model. The quest remains to identify a translational animal model of PD that optimally emulates the neuropathological and behavioral characteristics of PD with the timing of these deficits being protracted across the lifespan.

## Supporting information

Supplemental Results

## Acknowledgements

We thank the Division of Laboratory Animal Medicine at University of North Texas Health Science Center for exceptional animal care

## Declaration of interest

none.

## Ethics approval

All animals were used in compliance with the Office of Laboratory Animal Welfare guidelines and protocols approved by the Institutional Animal Care and Use Committee at the University of North Texas Health Science Center

## Consent for publication

Not applicable

## Availability of data and materials

Data generated or analyzed during this study are included in this published article and its supplementary information files. Additional data and materials are available from the corresponding author on reasonable request.

## Competing interests

The authors declare that they have no competing interests relevant to the content of this article, no financial or proprietary interests in any material discussed in this article, or affiliations with or involvement in any organization or entity with any financial interest or non-financial interest in the subject matter or materials discussed in this manuscript.

## Funding

This study was funded by the US Department of Defense U.S. Army Medical Research and Material Command Congressionally Directed Medical Research Program (Award Number W81XWH-19-1-0757) to MFS and Fellowship to IS by the National Institute on Aging Training Grant (Award Number T32AG020494). The funders had no role in the study design, in the collection, analysis, and interpretation of data, or writing of the manuscript.

## Authors’ contributions

Conceptualization: IS, MFS, VAN Data curation: IS, RM, WNB, KD, EAK. Formal analysis; IS, MFS, WNB, RM. Funding acquisition: MFS, IS, VAN. Project administration: IS, MFS, VAN. Interpretation of results: IS, MFS, VAN Writing original draft: IS, Review and editing: MFS, VAN, IS. All authors read and approved the final manuscript.

## Notes

### Competing Interest Statement

The authors have declared no competing interest.

## References

Aarsland, D., Creese, B., Politis, M., Chaudhuri, K.R., Ffytche, D.H., Weintraub, D., Ballard, C., 2017. Cognitive decline in Parkinson disease. Nat. Rev. Neurol. 13, 217–231.

Barker, R.A., Björklund, A., 2020. Animal models of Parkinson’s disease: Are they useful or not? J. Parkinsons. Dis. 10, 1335–1342.

Berg, D., Borghammer, P., Fereshtehnejad, S.-M., Heinzel, S., Horsager, J., Schaeffer, E., Postuma, R.B., 2021. Prodromal Parkinson disease subtypes - key to understanding heterogeneity. Nat. Rev. Neurol. 17, 349–361.

Bezard, E., Yue, Z., Kirik, D., Spillantini, M.G., 2013. Animal models of Parkinson’s disease: limits and relevance to neuroprotection studies. Mov. Disord. 28, 61–70.

Blackinton, J.G, Anuvert, A., Beilina, A., Olson, L., Cookson, M.R., Galter, D. 2007. Expression of Pink1 mRNA in human and rodent brain and in Parkinson’s disease. Brain Res. 1184, 10–16.

Bolam, J.P., Pissadaki, E.K., 2012. Living on the edge with too many mouths to feed: why dopamine neurons die. Mov. Disord. 27, 1478–1483.

Bonifati V, Rohe CF, Breedveld GJ, Fabrizio E, De Mari M, Tassorelli C, Tavella, et al. 2005. Early-onset parkinsonism associated with PINK1 mutations. Neurology 65: 87–95.

Borsche, M., König, I.R., Delcambre, S., Petrucci, S., Balck, A., Brüggemann, N., Zimprich, A., Wasner, K., Pereira, S.L., Avenali, M., Deuschle, C., Badanjak, K., Ghelfi, J., Gasser, T., Kasten, M., Rosenstiel, P., Lohmann, K., Brockmann, K., Valente, E.M., Youle, R.J., Grünewald, A., Klein, C., 2020. Mitochondrial damage-associated inflammation highlights biomarkers in PRKN/PINK1 parkinsonism. Brain 143, 3041–3051.

Braak, H., Del Tredici, K., Rüb, U., de Vos, R.A.I., Jansen Steur, E.N.H., Braak, E., 2003. Staging of brain pathology related to sporadic Parkinson’s disease. Neurobiol. Aging 24, 197– 211.

Brück, A., Aalto, S., Nurmi, E., Bergman, J., Rinne, J.O., 2005. Cortical 6-[18F]fluoro-L-dopa uptake and frontal cognitive functions in early Parkinson’s disease. Neurobiol. Aging 26, 891– 898.

Cai, X., Qiao, J., Knox, T., Iriah, S., Kulkarni, P., Madularu, D., Morrison, T., Waszczak, B., Hartner, J.C., Ferris, C.F., 2019. In search of early neuroradiological biomarkers for Parkinson’s Disease: Alterations in resting state functional connectivity and gray matter microarchitecture in PINK1 -/- rats. Brain Res. 1706, 58–67.

Cazeneuve, C., Sân, C., Ibrahim, S.A., Mukhtar, M.M., Kheir, M.M., Leguern, E., Brice, A., Salih, M.A., 2009. A new complex homozygous large rearrangement of the PINK1 gene in a Sudanese family with early onset Parkinson’s disease. Neurogenetics 10, 265–270.

Collier TJ, Kanaan NM, Kordower JH. 2011. Ageing as a primary risk factor for Parkinson’s disease: evidence from studies of non-human primates. Nat Rev Neurosci 12: 359–366.

Creed RB, Menalled L, Casey B, Dave KD, Janssens HB, Veinberg I, van der Hart M, Rassoulpour A, Goldberg MS. 2019. Basal and Evoked Neurotransmitter Levels in Parkin, DJ-1, PINK1 and LRRK2 Knockout Rat Striatum. Neuroscience 409, 169-179.

Creed, R.B., Roberts, R.C., Farmer, C.B., McMahon, L.L., Goldberg, M.S., 2021. Increased glutamate transmission onto dorsal striatum spiny projection neurons in Pink1 knockout rats. Neurobiol. Dis. 150, 105246.

Cullen, K.P., Grant, L.M., Kelm-Nelson, C.A., Brauer, A.F.L., Bickelhaupt, L.B., Russell, J.A., Ciucci, M.R., 2018. Pink1 -/- Rats Show Early-Onset Swallowing Deficits and Correlative Brainstem Pathology. Dysphagia 33, 749–758.

Dauer, W., Przedborski, S., 2003. Parkinson’s Disease: Mechanisms and Models. Neuron 39, 889–909.

de Haas, R., Heltzel, L.C.M.W., Tax, D., van den Broek, P., Steenbreker, H., Verheij, M.M.M., Russel, F.G.M., Orr, A.L., Nakamura, K., Smeitink, J.A.M., 2019. To be or not to be pink(1): contradictory findings in an animal model for Parkinson’s disease. Brain Commun 1, fcz016.

de Lau, L.M.L., Breteler, M.M.B., 2006. Epidemiology of Parkinson’s disease. Lancet Neurol. 5, 525–535.

DeAngelo, V.M., Hilliard, J.D., McConnell, G.C., 2022. Dopaminergic but not cholinergic neurodegeneration is correlated with gait disturbances in PINK1 knockout rats. Behav. Brain Res. 417, 113575.

Dirnberger, G., Jahanshahi, M., 2013. Executive dysfunction in Parkinson’s disease: a review. J. Neuropsychol. 7, 193–224.

Durcan, R., Wiblin, L., Lawson, R.A., Khoo, T.K., Yarnall, A.J., Duncan, G.W., Brooks, D.J., Pavese, N., Burn, D.J., ICICLE-PD Study Group, 2019. Prevalence and duration of non-motor symptoms in prodromal Parkinson’s disease. Eur. J. Neurol. 26, 979–985.

Ennaceur, A., 2010. One-trial object recognition in rats and mice: methodological and theoretical issues. Behav. Brain Res. 215, 244–254.

Fallon, S.J., Williams-Gray, C.H., Barker, R.A., Owen, A.M., Hampshire, A., 2013. Prefrontal dopamine levels determine the balance between cognitive stability and flexibility. Cereb. Cortex 23, 361–369.

Ferris, C.F., Morrison, T.R., Iriah, S., Malmberg, S., Kulkarni, P., Hartner, J.C., Trivedi, M., 2018. Evidence of Neurobiological Changes in the Presymptomatic PINK1 Knockout Rat. J. Parkinsons. Dis. 8, 281–301.

Gelders, G., Baekelandt, V., Van der Perren, A., 2018. Linking Neuroinflammation and Neurodegeneration in Parkinson’s Disease. J Immunol Res 2018, 4784268.

González-Rodríguez, P., Zampese, E., Stout, K.A., Guzman, J.N., et al. 2021. Disruption of mitochondrial complex I induces progressive parkinsonism. Nature 599, 650–6.

Grant, L.M., Kelm-Nelson, C.A., Hilby, B.L., Blue, K.V., Paul Rajamanickam, E.S., Pultorak, J.D., Fleming, S.M., Ciucci, M.R., 2015. Evidence for early and progressive ultrasonic vocalization and oromotor deficits in a PINK1 gene knockout rat model of Parkinson’s disease. J. Neurosci. Res. 93, 1713–1727.

Grigoruţă, M., Martínez-Martínez, A., Dagda, R.Y., Dagda, R.K., 2020. Psychological Stress Phenocopies Brain Mitochondrial Dysfunction and Motor Deficits as Observed in a Parkinsonian Rat Model. Mol. Neurobiol. 57, 1781–1798.

Haddad, D., Nakamura, K., 2015. Understanding the susceptibility of dopamine neurons to mitochondrial stressors in Parkinson’s disease. FEBS Lett. 589, 3702–3713.

Hall, S., Janelidze, S., Surova, Y., Widner, H., Zetterberg, H., Hansson, O., 2018. Cerebrospinal fluid concentrations of inflammatory markers in Parkinson’s disease and atypical parkinsonian disorders. Sci. Rep. 8, 13276.

Hartmann, A., 2004. Postmortem studies in Parkinson’s disease. Dialogues Clin. Neurosci. 6, 281–293.

Hauser, R.A., 2018. Help cure Parkinson’s disease: please don’t waste the Golden Year. NPJ Parkinsons Dis 4, 29.

Hayashida A., Li Y, Yoshino H, Daida K, Ikeda A, Ogaki K, Fuse A, Mori A, Takanashi M, et al., 2021. The identified clinical features of Parkinson’s disease in homo-, heterozygous and digenic variants of PINK1. Neurobiol Aging 97: 146e1–146e13

Heng, N., Malek, N., Lawton, M.A., Nodehi, A., Pitz, V., Grosset, K.A., Ben-Shlomo, Y., Grosset, D.G., 2023. Striatal Dopamine Loss in Early Parkinson’s Disease: Systematic Review and Novel Analysis of Dopamine Transporter Imaging. Mov Disord Clin Pract 10, 539–546.

Hindle, J.V., 2010. Ageing, neurodegeneration and Parkinson’s disease. Age Ageing 39, 156– 161.

Hirsch, E.C., Breidert, T., Rousselet, E., Hunot, S., Hartmann, A., Michel, P.P., 2003. The role of glial reaction and inflammation in Parkinson’s disease. Ann. N. Y. Acad. Sci. 991, 214–228.

Hoffmeister, J., 2021. The Role of Norepinephrine in Vocal Communication and Anxiety in Parkinson Disease. University of Wisconsin--Madison.

Hoffmeister, J.D., Kelm-Nelson, C.A., Ciucci, M.R., 2021. Quantification of brainstem norepinephrine relative to vocal impairment and anxiety in the Pink1-/- rat model of Parkinson disease. Behav. Brain Res. 414, 113514.

Hoffmeister, J.D., Kelm-Nelson, C.A., Ciucci, M.R., 2022. Manipulation of vocal communication and anxiety through pharmacologic modulation of norepinephrine in the Pink1-/- rat model of Parkinson disease. Behav. Brain Res. 418, 113642.

Ishihara-Paul L, Hulihan MM, Kachergus J, Upmanyu R, Warren L, Amouri R, Elango R et al. 2008. Pink1 mutations and parkinsonism. Neurology 71: 896–902.

Kaasinen, V., Nurmi, E., Brück, A., Eskola, O., Bergman, J., Solin, O., Rinne, J.O., 2001. Increased frontal [18F]fluorodopa uptake in early Parkinson’s disease: sex differences in the prefrontal cortex. Brain 124, 1125–1130.

Kasanga, E.A., Han, Y., Shifflet, M.K., Navarrete, W., McManus, R., Parry, C., Barahona, A., Nejtek, V.A., Manfredsson, F.P., Kordower, J.H., Richardson, J.R., Salvatore, M.F., 2023. Nigral-specific increase in ser31 phosphorylation compensates for tyrosine hydroxylase protein and nigrostriatal neuron loss: Implications for delaying parkinsonian signs. Exp. Neurol. 368, 114509.

Kasten, M., Hartmann, C., Hampf, J., Schaake, S., Westenberger, A., Vollstedt, E.-J., Balck, A., Domingo, A., Vulinovic, F., Dulovic, M., Zorn, I., Madoev, H., Zehnle, H., Lembeck, C.M., Schawe, L., Reginold, J., Huang, J., König, I.R., Bertram, L., Marras, C., Lohmann, K., Lill, C.M., Klein, C., 2018. Genotype-phenotype relations for the Parkinson’s disease genes Parkin, PINK1, DJ1: MDSGene systematic review. Mov. Disord. 33, 730–741.

Kelm-Nelson, C.A., Lechner, S.A., Samantha, E., Kaldenberg, T.A.R., Natalie, K., Ciucci, M.R., 2021. Pink1−/− rats are a useful tool to study early Parkinson disease. Brain Commun 3, fcab077.

Kelm-Nelson, C.A., Trevino, M.A., Ciucci, M.R., 2018. Quantitative analysis of catecholamines in the Pink1 -/- rat model of early-onset Parkinson’s disease. Neuroscience 379, 126–141.

Khoo, S.Y.-S., Uhrig, A., Samaha, A.-N., Chaudhri, N., 2022. Does vendor breeding colony influence sign- and goal-tracking in Pavlovian conditioned approach? A preregistered empirical replication. Neuroanat. Behav. 4, e46.

Kitada, T., Pisani, A., Porter, D. R., Yamaguchi, H., Tscherter, A., Martella, G., Bonsi, P., Zhang, C., Pothos, E. N., & Shen, J. 2007. Impaired dopamine release and synaptic plasticity in the striatum of PINK1-deficient mice. Proc Natl Acad Sci USA (27), 11441–11446. 10.1073/pnas.0702717104

Kitada, T., Tong, Y., Gautier, C. A., & Shen, J. 2009. Absence of nigral degeneration in aged parkin/DJ-1/PINK1 triple knockout mice. J Neurochem 111(3), 696–702. 10.1111/j.1471-4159.2009.06350.x

Klein, C., Lohmann-Hedrich, K., Rogaeva, E., Schlossmacher, M.G., Lang, A.E., 2007. Deciphering the role of heterozygous mutations in genes associated with parkinsonism. Lancet Neurol. 6, 652–662.

Kordower, J.H., Olanow, C.W., Dodiya, H.B., Chu, Y., Beach, T.G., Adler, C.H., Halliday, G.M., Bartus, R.T., 2013. Disease duration and the integrity of the nigrostriatal system in Parkinson’s disease. Brain 136, 2419–2431.

Kumar, A., Tamjar, J., Waddell, A.D., Woodroof, H.I., Raimi, O.G., Shaw, A.M., Peggie, M., Muqit, M.M., van Aalten, D.M., 2017. Structure of PINK1 and mechanisms of Parkinson’s disease-associated mutations. Elife 6. 10.7554/eLife.29985

Lechner SA, Welsch JM, Phapill NK, Kaldenberg TAR, Regenbaum A, Kelm-Nelson CA. 2022. Predictors of prodromal Parkinson’s disease in young adult Pink1−/− rats. Front Behav Neurosci. 2022;16.

Ma, K.Y., Fokkens, M.R., van Laar, T., Verbeek, D.S., 2021. Systematic analysis of PINK1 variants of unknown significance shows intact mitophagy function for most variants. NPJ Parkinsons Dis 7, 113.

Marquis, J.M., Lettenberger, S.E., Kelm-Nelson, C.A., 2020. Early-onset Parkinsonian behaviors in female Pink1-/- rats. Behav. Brain Res. 377, 112175.

Marras, C., Beck, J.C., Bower, J.H., Roberts, E., Ritz, B., Ross, G.W., Abbott, R.D., Savica, R., Van Den Eeden, S.K., Willis, A.W., Tanner, C.M., Parkinson’s Foundation P4 Group, 2018. Prevalence of Parkinson’s disease across North America. NPJ Parkinsons Dis 4, 21.

Maynard, M.E., Redell, J.B., Kobori, N., Underwood, E.L., Fischer, T.D., Hood, K.N., LaRoche, V., Waxham, M.N., Moore, A.N., Dash, P.K., 2020. Loss of PTEN-induced kinase 1 (Pink1) reduces hippocampal tyrosine hydroxylase and impairs learning and memory. Exp. Neurol. 323, 113081.

Mazumder, S., Bahar, A.Y., Shepherd, C.E., Prasad, A.A., 2022. Post-mortem brain histological examination in the substantia nigra and subthalamic nucleus in Parkinson’s disease following deep brain stimulation. Front. Neurosci. 16, 948523.

McDonald, C., Gordon, G., Hand, A., Walker, R.W., Fisher, J.M., 2018. 200 Years of Parkinson’s disease: what have we learnt from James Parkinson? Age Ageing 47, 209–214.

McNaught, K.S.P., Olanow, C.W., Halliwell, B., Isacson, O., Jenner, P., 2001. Failure of the ubiquitin–proteasome system in Parkinson’s disease. Nat. Rev. Neurosci. 2, 589–594.

Murphy, B.L., Arnsten, A.F., Goldman-Rakic, P.S., Roth, R.H., 1996. Increased dopamine turnover in the prefrontal cortex impairs spatial working memory performance in rats and monkeys. Proc. Natl. Acad. Sci. U. S. A. 93, 1325–1329.

Nejtek VA, James RN, Salvatore MF, Alphonso HM, Boehm GW. 2021. Premature cognitive decline in specific domains found in young veterans with mTBI coincide with elder normative scores and advanced-age subjects with early-stage Parkinson’s disease. PLoS ONE 16: e0258851.

Nybø, C.J., Gustavsson, E.K., Farrer, M.J., Aasly, J.O., 2020. Neuropathological findings in PINK1-associated Parkinson’s disease. Parkinsonism Relat. Disord. 78, 105–108.

Oliff, H.S., Weber, E., Eilon, G., Marek, P., 1995. The role of strain/vendor differences on the outcome of focal ischemia induced by intraluminal middle cerebral artery occlusion in the rat. Brain Res. 675, 20–26.

Pinizzotto, C.C., Dreyer, K.M., Aje, O.A., Caffrey, R.M., Madhira, K., Kritzer, M.F., 2022. Spontaneous Object Exploration in a Recessive Gene Knockout Model of Parkinson’s Disease: Development and Progression of Object Recognition Memory Deficits in Male Pink1–/– Rats. Front. Behav. Neurosci. 16. 10.3389/fnbeh.2022.951268

Polinski, N.K., 2021. A summary of phenotypes observed in the *in vivo* rodent alpha-synuclein preformed fibril model. J Parkinsons Dis. 11, 1555–1567.

Postuma, R.B., Berg, D., 2019. Prodromal Parkinson’s Disease: The Decade Past, the Decade to Come. Mov. Disord. 34, 665–675.

Pruett, B.S., Salvatore, M.F., 2013. Nigral GFRα1 infusion in aged rats increases locomotor activity, nigral tyrosine hydroxylase, and dopamine content in synchronicity. Mol Neurobiol 47, 988–999.

Quinn, R., 2005. Comparing rat’s to human’s age: how old is my rat in people years? Nutrition 21, 775–777.

Ren, X., Butterfield, D.A., 2021. Fidelity of the PINK1 knockout rat to oxidative stress and other characteristics of Parkinson disease. Free Radic. Biol. Med. 163, 88–101.

Ricciardi L, Petrucci S, Guidubaldi A, Ialongo T, Serra L, Ferraris A, Spano B, Bozzali M, Valente EM, Bentivoglio AR. 2014. Phenotypic variability of Pink1 expression: 12 years’ clinical follow-up of two Italian families. Mov Disord 29: 1561–1566.

Salvatore, M.F., Terrebonne, J., Fields, V., Nodurft, D., Runfalo, C., Lattimer, B., Ingram, D.K. 2016b. Initiation of calorie restriction in middle-aged male rats attenuates aging-related motoric decline and bradykinesia without increased striatal dopamine. Neurobiol. Aging 37, 192–207.

Salvatore, M.F., Terrebonne, J., Cantu, M.A., McInnis, T.R., Venable, K., Kelley, P., Kasanga, E.A., Latimer, B., Owens, C.L., et al. 2017. Dissociation of striatal dopamine and tyrosine hydroxylase expression in aging-related motor decline: evidence from calorie restriction intervention. J. Gerontol. Biol. Sci. 73, 11–20.

Salvatore, M.F. McInnis, T.R., Cantu, M.A., Apple, D.M., Pruett, B.S. 2019. Tyrosine hydroxylase inhibition in substantia nigra decreases movement frequency. Mol Neurobiol. 56, 2728–2740.

Salvatore, M.F., Soto, I., Alphonso, H., Cunningham, R., James, R., Nejtek, V.A., 2021. Is there a Neurobiological Rationale for the Utility of the Iowa Gambling Task in Parkinson’s Disease? J. Parkinsons. Dis. 11, 405–419.

Salvatore, M.F., Soto, I., Kasanga, E.A., James, R., Shifflet, M.K., Doshier, K., Little, J.T., John, J., Alphonso, H.M., Cunningham, J.T., Nejtek, V.A., 2022. Establishing Equivalent Aerobic Exercise Parameters Between Early-Stage Parkinson’s Disease and Pink1 Knockout Rats. J. Parkinsons. Dis. 12, 1897–1915.

Salvatore, M.F., Kasanga, E.A., Kelly, D.P., Venable, K.E., McInnis, T.R., Cantu, M.A., Terrebonne, J., Lanza, K., Meadows, S.M., Centner, A., Bishop, C., Ingram, D.K. 2023. Modulation of nigral dopamine signaling mitigates parkinsonian signs of aging: evidence from intervention with caloric restriction or inhibition of dopamine uptake. GeroScience 45, 45–63.

Salvatore, M.F. 2024. Dopamine Signaling in Substantia Nigra and Its Impact on Locomotor Function-Not a New Concept, but Neglected Reality. Int J Mol Sci. 25(2). doi:10.3390/ijms25021131

Schubert, A.F., Gladkova, C., Pardon, E., Wagstaff, J.L., Freund, S.M.V., Steyaert, J., Maslen, S.L., Komander, D., 2017. Structure of PINK1 in complex with its substrate ubiquitin. Nature 552, 51–56.

Soto, I., Nejtek, V.A., Siderovski, DP, Salvatore, M.F. 2024. PINK1 knockout rats show premotor cognitive deficits measured through a complex maze. bioRxiv. doi:10.1101/2024.01.18.576285

Steele, J.C., Guella, I., Szu-Tu, C., Lin, M.K., Thompson, C., Evans, D.M., Sherman, H.E., Vilariño-Güell, C., Gwinn, K., Morris, H., Dickson, D.W., Farrer, M.J., 2015. Defining neurodegeneration on Guam by targeted genomic sequencing. Ann. Neurol. 77, 458–468.

Stefanis, L., 2012. α-Synuclein in Parkinson’s disease. Cold Spring Harb. Perspect. Med. 2, a009399.

Stoker, T.B., Greenland, J.C., 2018. Parkinson’s Disease: Pathogenesis and Clinical Aspects. Codon Publications.

Sun, J., Kouranova, E., Cui, X., Mach, R.H., Xu, J., 2013. Regulation of dopamine presynaptic markers and receptors in the striatum of DJ-1 and Pink1 knockout rats. Neurosci. Lett. 557 Pt B, 123–128.

Taghavi, S., Chaouni, R., Tafakhori, A., Azcona, L.J., Firouzabadi, S.G., Omrani, M.D., Jamshidi, J., Emamalizadeh, B., Shahidi, G.A., Ahmadi, M., Habibi, S.A.H., Ahmadifard, A., Fazeli, A., Motallebi, M., Petramfar, P., Askarpour, S., Askarpour, S., Shahmohammadibeni, H.A., Shahmohammadibeni, N., Eftekhari, H., Shafiei Zarneh, A.E., Mohammadihosseinabad, S., Khorrami, M., Najmi, S., Chitsaz, A., Shokraeian, P., Ehsanbakhsh, H., Rezaeidian, J., Ebrahimi Rad, R., Madadi, F., Andarva, M., Alehabib, E., Atakhorrami, M., Mortazavi, S.E., Azimzadeh, Z., Bayat, M., Besharati, A.M., Harati-Ghavi, M.A., Omidvari, S., Dehghani-Tafti, Z., Mohammadi, F., Mohammad Hossein Pour, B., Noorollahi Moghaddam, H., Esmaili Shandiz, E., Habibi, A., Taherian-Esfahani, Z., Darvish, H., Paisán-Ruiz, C., 2018. A Clinical and Molecular Genetic Study of 50 Families with Autosomal Recessive Parkinsonism Revealed Known and Novel Gene Mutations. Mol. Neurobiol. 55, 3477–3489.

Urrutia, P.J., Mena, N.P., Núñez, M.T., 2014. The interplay between iron accumulation, mitochondrial dysfunction, and inflammation during the execution step of neurodegenerative disorders. Front. Pharmacol. 5, 38.

Valente, E.M., Abou-Sleiman, P.M., Caputo, V., Muqit, M.M.K., Harvey, K., Gispert, S., Ali, Z., Del Turco, D., Bentivoglio, A.R., Healy, D.G., Albanese, A., Nussbaum, R., González-Maldonado, R., Deller, T., Salvi, S., Cortelli, P., Gilks, W.P., Latchman, D.S., Harvey, R.J., Dallapiccola, B., Auburger, G., Wood, N.W., 2004. Hereditary early-onset Parkinson’s disease caused by mutations in PINK1. Science 304, 1158–1160.

Villeneuve, L.M., Purnell, P.R., Boska, M.D., Fox, H.S., 2016. Early Expression of Parkinson’s Disease-Related Mitochondrial Abnormalities in PINK1 Knockout Rats. Mol. Neurobiol. 53, 171–186.

Wood-Kaczmar, A., Gandhi, S., Yao, Z., Abramov, A.Y., Miljan, E.A., Keen, G., Stanyer, L., Hargreaves, I., Klupsch, K., Deas, E., Downward, J., Mansfield, L., Jat, P., Taylor, J., Heales, S., Duchen, M.R., Latchman, D., Tabrizi, S.J., Wood, N.W., 2008. PINK1 is necessary for long term survival and mitochondrial function in human dopaminergic neurons. PLoS One 3, e2455.

Zeiss, C.J., Allore, H.G., Beck, A.P., 2017. Established patterns of animal study design undermine translation of disease-modifying therapies for Parkinson’s disease. PLoS One 12, e0171790.

Zhang, T.D., Kolbe, S.C., Beauchamp, L.C., Woodbridge, E.K., Finkelstein, D.I., Burrows, E.L., 2022. How Well Do Rodent Models of Parkinson’s Disease Recapitulate Early Non-Motor Phenotypes? A Systematic Review. Biomedicines 10. 10.3390/biomedicines10123026

Zhi, L., Qin, Q., Muqeem, T., Seifert, E. L., Liu, W., Zheng, S., Li, C., & Zhang, H. 2019. Loss of PINK1 causes age-dependent decrease of dopamine release and mitochondrial dysfunction. Neurobiol Aging, 75, 1–10. 10.1016/j.neurobiolaging.2018.10.025

