## Supplemental Results for "Aging hastens locomotor decline in PINK1 knockout rats in association with decreased nigral, but not striatal, dopamine and tyrosine hydroxylase expression"

A.
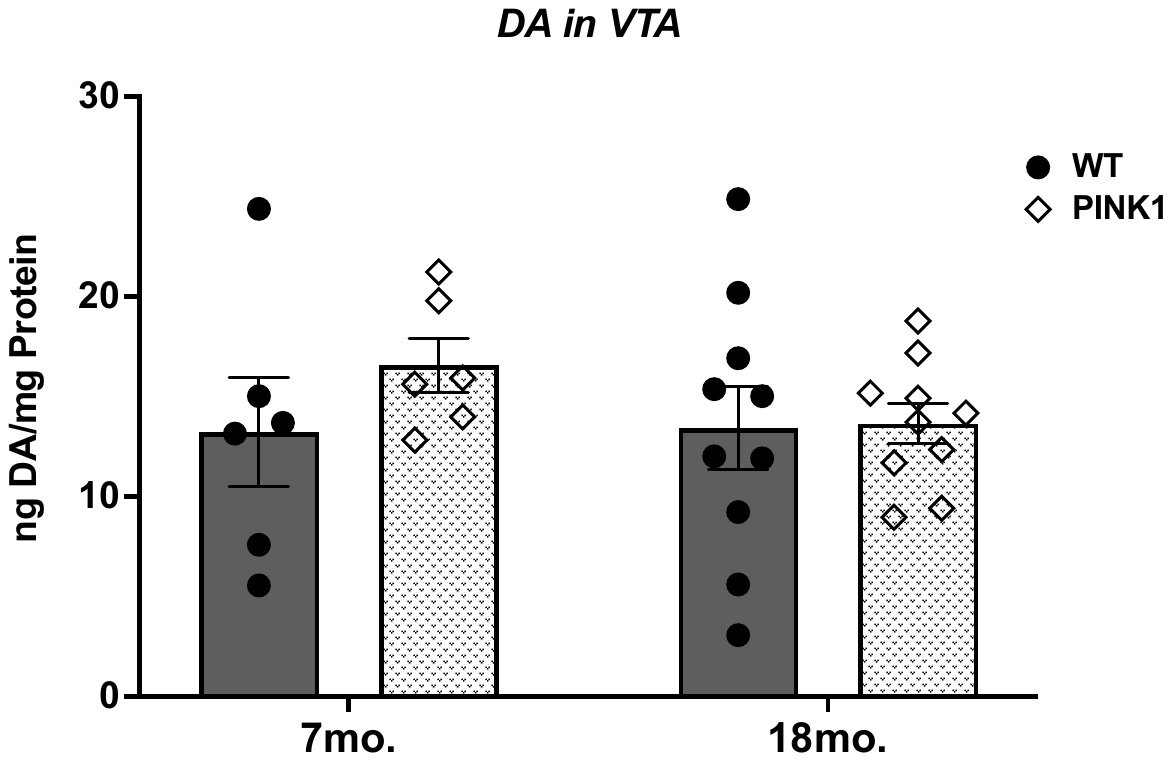

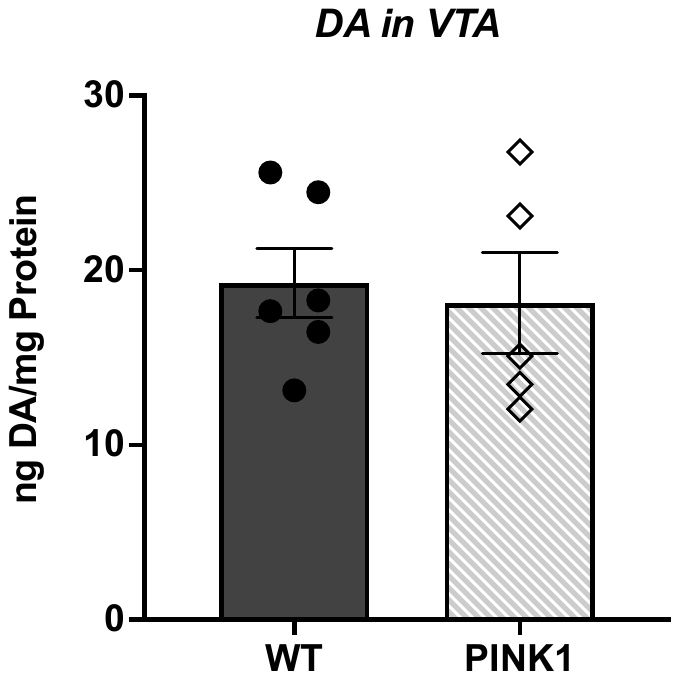
 **B*.***

***Supplemental Figure 1: DA in VTA***

There was no significant difference in DA content in the VTA **(A)** between16-month-old KO rats compared to age-matched WT rats (t= 0.08, ns, df= 10)**.** Similarly**,** in the second cohort **(B),** there was no age or genotype effect in DA content in the VTA. 7mo. WT vs 7mo. KO (t= 1.12, ns, df= 28); 7mo. WT vs 18mo. WT (t= 0.07, ns, df= 28); 7mo. KO vs 18mo. KO (t= 1.10, ns, df= 28); 18mo. WT vs 18mo. KO (t= 0.09, ns, df= 28).

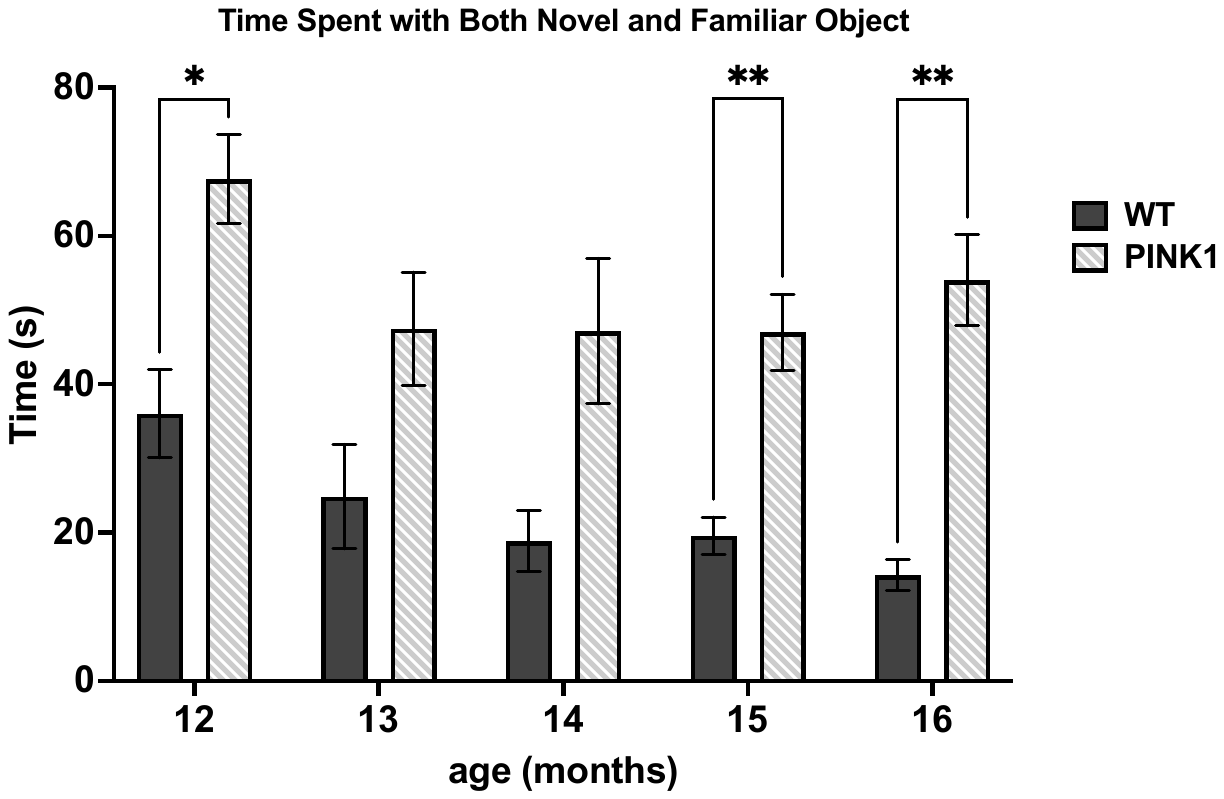
***Supplemental Figure 2: Total Time Spent with Objects in NOR testing.***

The PINK1 KO rats spent more time interacting with both the novel and familiar objects, particularly starting at 12mo. and onward. Age (F(4,40)= 2.58, ns). Genotype (F(1,11)= 65.29, *p<*0.0001) Age x Genotype (F(4,40)= 0.76, ns). WT vs KO 12mo. (t= 5.15, **p*=0.016, df= 51); 13mo. (t= 2.18, ns, df= 51); 14mo. (t= 2.20, ns, df= 51); 15mo. (t=4.01, ***p*= 0.008, df= 51); 16mo. (t= 5.28, ***p*= 0.004, df= 51).
